# *Drosophila* Evolution over Space and Time (DEST) - A New Population Genomics Resource

**DOI:** 10.1101/2021.02.01.428994

**Authors:** Martin Kapun, Joaquin C. B. Nunez, María Bogaerts-Márquez, Jesús Murga-Moreno, Margot Paris, Joseph Outten, Marta Coronado-Zamora, Courtney Tern, Omar Rota-Stabelli, Maria P. García Guerreiro, Sònia Casillas, Dorcas J. Orengo, Eva Puerma, Maaria Kankare, Lino Ometto, Volker Loeschcke, Banu S. Onder, Jessica K. Abbott, Stephen W. Schaeffer, Subhash Rajpurohit, Emily L Behrman, Mads F. Schou, Thomas J.S. Merritt, Brian P Lazzaro, Amanda Glaser-Schmitt, Eliza Argyridou, Fabian Staubach, Yun Wang, Eran Tauber, Svitlana V. Serga, Daniel K. Fabian, Kelly A. Dyer, Christopher W. Wheat, John Parsch, Sonja Grath, Marija Savic Veselinovic, Marina Stamenkovic-Radak, Mihailo Jelic, Antonio J. Buendía-Ruíz, M. Josefa Gómez-Julián, M. Luisa Espinosa-Jimenez, Francisco D. Gallardo-Jiménez, Aleksandra Patenkovic, Katarina Eric, Marija Tanaskovic, Anna Ullastres, Lain Guio, Miriam Merenciano, Sara Guirao-Rico, Vivien Horváth, Darren J. Obbard, Elena Pasyukova, Vladimir E. Alatortsev, Cristina P. Vieira, Jorge Vieira, J. Roberto Torres, Iryna Kozeretska, Oleksandr M. Maistrenko, Catherine Montchamp-Moreau, Dmitry V. Mukha, Heather E. Machado, Antonio Barbadilla, Dmitri Petrov, Paul Schmidt, Josefa Gonzalez, Thomas Flatt, Alan O. Bergland

## Abstract

*Drosophila melanogaster* is a leading model in population genetics and genomics, and a growing number of whole-genome datasets from natural populations of this species have been published over the last 20 years. A major challenge is the integration of these disparate datasets, often generated using different sequencing technologies and bioinformatic pipelines, which hampers our ability to address questions about the evolution and population structure of this species. Here we address these issues by developing a bioinformatics pipeline that maps pooled sequencing (Pool-Seq) reads from *D. melanogaster* to a hologenome consisting of fly and symbiont genomes and estimates allele frequencies using either a heuristic (PoolSNP) or a probabilistic variant caller (SNAPE-pooled). We use this pipeline to generate the largest data repository of genomic data available for *D. melanogaster* to date, encompassing 271 population samples from over 100 locations in >20 countries on four continents based on a combination of 121 unpublished and 150 previously published genomic datasets. Several of these locations have been sampled at different seasons across multiple years. This dataset, which we call *Drosophila Evolution over Space and Time* (DEST), is coupled with sampling and environmental meta-data. A web-based genome browser and web portal provide easy access to the SNP dataset. Our aim is to provide this scalable platform as a community resource which can be easily extended via future efforts for an even more extensive cosmopolitan dataset. Our resource will enable population geneticists to analyze spatio-temporal genetic patterns and evolutionary dynamics of *D. melanogaster* populations in unprecedented detail.

## Introduction

The vinegar fly *Drosophila melanogaster* is one of the oldest and most important genetic model systems and has played a key role in the development of theoretical and empirical population genetics (e.g., Schneider 2000; Larracuente and Roberts 2015; Haudry *et al*. 2020). Through decades of work, we now have a basic picture of the evolutionary origin (David and Capy 1988; Lachaise *et al*. 1988; Keller 2007; Sprengelmeyer *et al*. 2020), colonization history and demography (Caracristi and Schlötterer 2003; Li and Stephan 2006; Duchen *et al*. 2013; Grenier *et al*. 2015; Arguello *et al*. 2019; Kapopoulou *et al*. 2020), and spatio-temporal diversification patterns of this species and its close relatives (Kolaczkowski *et al*. 2011; Fabian *et al*. 2012; Bergland *et al*. 2014; Lack *et al*. 2016; Machado *et al*. 2016; Kapun *et al*. 2016, 2020). The availability of high-quality reference genomes (Adams 2000; Celniker and Rubin 2003; dos Santos *et al*. 2015) and genetic tools (Schneider 2000; Duffy 2002; Jennings 2011; Hales *et al*. 2015; Haudry *et al*. 2020) facilitates placing evolutionary studies of flies in a mechanistic context, allowing for the functional characterization of ecologically relevant polymorphisms (e.g., de Jong and Bochdanovits 2003; Paaby *et al*. 2010, 2014; Mateo *et al*. 2014; Kapun *et al*. 2016; Durmaz *et al*. 2018, 2019; Ramaekers *et al*. 2019).

Recently, work on the evolutionary biology of *Drosophila* has been fueled by a growing number of population genomic datasets from field collections across a large portion of *D. melanogaster*’s range (Grenier *et al*. 2015; Machado et al. 2021; Guirao-Rico and González 2019; Arguello *et al*. 2019). These genomic data consist either of re-sequenced inbred (or haploid) individuals (e.g., Mackay *et al*. 2012; Langley *et al*. 2012; Grenier *et al*. 2015; Lack *et al*. 2015, 2016; Mateo *et al*. 2018; Kapopoulou *et al*. 2020) or pooled sequencing of outbred population samples (Pool-Seq; e.g., Kolaczkowski *et al*. 2011; Fabian *et al*. 2012; Bastide *et al*. 2013; Campo *et al*. 2013; Bergland *et al*. 2014; Machado *et al*. 2016, 2019; Kapun *et al*. 2016, 2020). Pooled re-sequencing provides accurate and precise estimates of allele frequencies across most of the allele frequency spectrum (Zhu *et al*. 2012; Lynch *et al*. 2014; Schlötterer *et al*. 2014) at a fraction of the cost of individual-based sequencing. Although Pool-Seq retains limited information about linkage disequilibrium (LD) relative to individual sequencing (Feder *et al*. 2012), Pool-Seq data can be used to infer complex demographic histories (e.g., Cheng *et al*. 2012; Bergland *et al*. 2016; Deitz *et al*. 2016; Gould *et al*. 2017; Corbett-Detig and Nielsen 2017; Giesen *et al*. 2020), characterize levels of diversity (Kofler *et al*. 2011a, 2011b; Ferretti *et al*. 2013; Kapun *et al*. 2020), and infer genomic loci involved in recent adaptation in nature (Flatt 2016; Kapun *et al*. 2016, 2020; Gould *et al*. 2017; Bogaerts-Márquez *et al*. 2020; Machado et al. 2021) and during experimental evolution (e.g., Turner *et al*. 2011; Orozco-terWengel *et al*. 2012; Burke 2012; Kofler and Schlötterer 2014). However, the rapidly increasing number of genomic datasets processed with different bioinformatic pipelines makes it difficult to compare results across studies and to jointly analyze multiple datasets. Differences among bioinformatic pipelines include filtering methods for the raw reads, mapping algorithms, the choice of the reference genome or SNP calling approaches, potentially generating biases when combining processed datasets from different sources for joint analyses (e.g., Gautier *et al*. 2013; Hoban *et al*. 2016).

To address these issues, we have developed a modular bioinformatics pipeline to map Pool-Seq reads to a hologenome consisting of fly and microbial genomes, to remove reads from potential *Drosophila simulans* contaminants, and to estimate allele frequencies using two complementary SNP callers. Our pipeline is available as a Docker image (available from https://dest.bio) to standardize versions of software used for filtering and mapping, to make the pipeline available independently of the operating system used, and to facilitate future updates and modification of the pipeline. In addition, our pipeline allows using either heuristic or probabilistic methods for SNP calling, based on PoolSNP (Kapun *et al*. 2020) and SNAPE-pooled (Raineri *et al*. 2012). We also provide tools for performing *in-silico* pooling of existing inbred (haploid) lines that exist as part of other *Drosophila* population genomic resources (Pool *et al*. 2012; Langley *et al*. 2012; Grenier *et al*. 2015; Kao *et al*. 2015; Lack *et al*. 2015, 2016). This pipeline is also designed to be flexible, facilitating the streamlined addition of new population samples as they arise.

Using this pipeline, we generated a unified dataset of pooled allele frequency estimates of *D. melanogaster* sampled across a large portion of its world-wide distribution, including Europe, North America, Africa, Australia, and Asia. This dataset is the result of the collaborative efforts of the European DrosEU (Kapun *et al*. 2020) and DrosRTEC (Machado et al. 2021) consortia and combines both novel and previously published population genomic data. Our dataset combines samples from 100 localities, 55 of which were sampled at two or more time points across the reproductive season (~10-15 generations/year) for one or more years. Collectively, these samples represent >13,000 individuals, cumulatively sequenced to >16,000x coverage or ~1x per fly. The cost-effectiveness of Pool-Seq has enabled us to estimate genome-wide allele frequencies over geographic space (continental and sub-continental) and time (seasonal, annual and decadal) scales, thus making our data a unique resource for advancing our understanding of fundamental adaptive and neutral evolutionary processes. We provide data in two file formats (VCF and GDS: Danecek *et al*. 2011; Zheng *et al*. 2017), thus allowing researchers to utilize a variety of tools for computational analyses. Our dataset also contains sampling and environmental meta-data to enable various downstream analyses of biological interest.

## Materials and Methods

### Data sources

The genomic dataset presented here has been assembled from a combination of Pool-Seq libraries and *in-silico* pooled haplotypes. We combined 246 Pool-Seq libraries of population samples from Europe, North America and the Caribbean that were sampled through space and time by two collaborating consortia in North America (DrosRTEC: https://web.sas.upenn.edu/paul-schmidt-lab/dros-rtec/) and Europe (DrosEU: http://droseu.net) between 2003 and 2016. Of these 246 Pool-Seq samples, 121 samples represent previously unpublished samples generated by DrosEU, 48 DrosEU samples previously reported in Kapun *et al*. (2020), and 77 samples previously reported in Machado *et al*. (2021). In addition, we integrated genomic data from >900 inbred or haploid genomes from 25 populations in Africa, Europe, Australia, and North America available from the *Drosophila* Genome Nexus dataset (DGN v1.1; Pool *et al*. 2012; Langley *et al*. 2012; Grenier *et al*. 2015; Kao *et al*. 2015; Lack *et al*. 2015, 2016) We further included the *D. simulans* haplotype (w^501^; Hu *et al*. 2013), built as part of the DGN dataset, as an outgroup, making this repository of 272 (246 Pool-Seq + 25 DGN + 1 *D. simulans*) whole-genome sequenced samples the largest dataset of genome-wide SNP polymorphisms available for *D. melanogaster* to date.

### Metadata

We assembled uniform meta-data for all samples (Supplementary Material online, supplementary table S1). This information includes collection coordinates, collection date, and the number of flies per sample. Samples are also linked to bioclimatic variables from the nearest WorldClim (Hijmans *et al*. 2005) raster cell at a resolution of 2.5° and to weather stations from the Global Historical Climatology Network (GHCND; ftp://ftp.ncdc.noaa.gov/pub/data/ghcn/daily/) to allow for future analyses of the environmental drivers that might underlie genetic change. We also provide summaries of basic attributes of each sample derived from the sequencing data including average read depth, PCR duplicate rate, *D. simulans* contamination rate, relative abundances of non-synonymous versus synonymous polymorphisms (*p*_N_/*p*_S_), the number of private polymorphisms, diversity statistics (Watterson’s *θ, π* and Tajima’s *D*), and estimates of inversion frequencies.

### Sample collection

The majority of population samples contributed by the DrosEU and the DrosRTEC consortia was collected in a coordinated fashion to generate a consistent dataset with minimized sampling bias. In brief, fly collections were performed exclusively in natural or semi-natural habitats, such as orchards, vineyards and compost piles. For most European collections, flies were collected using mashed banana, or apples with live yeast as bait in traps placed at sampling sites for multiple days to attract flies, or by sweep netting (see Kapun *et al*. 2020 for more details). For North American collections, flies were collected by sweep-net, aspiration, or baiting over natural substrate or using baited traps (see Behrman *et al*. 2018; Machado et al. 2021 for details). Samples were either field-caught flies (n=227), from F1 offspring of wild-caught females (n=7), from a mixture of F1 and wild-caught flies (n=7), or from flies kept as isofemale lines in the lab for 5 generations or less (n=4); see supplementary table 1 for more information. To minimize cross-contamination with the closely related sympatric sister species *D. simulans*, we only sequenced male *D. melanogaster* specimens, allowing for higher confidence discrimination between the two species based on the morphology of male genitalia (Capy and Gibert 2004; Markow and O’Grady 2006). Samples were stored in 95% ethanol at −20°C before DNA extraction.

### DNA extraction and sequencing

The DrosEU and DrosRTEC consortia centralized extractions from pools of flies. DNA was extracted either using chloroform/phenol-based (DrosEU: Kapun *et al*. 2020) or lithium chloride/potassium acetate extraction protocols (DrosRTEC: Bergland *et al*. 2014; Machado et al. 2021) after homogenization with bead beating or a motorized pestle. DrosEU samples from the 2014 collection were sequenced on an Illumina NextSeq 500 sequencer at the Genomics Core Facility of Pompeu Fabra University in Barcelona, Spain. Libraries of the previously unpublished DrosEU samples from 2015 and 2016 were constructed using the Illumina TruSeq PCR Free library preparation kit following the manufacturer’s instructions and sequenced on the Illumina HiSeq X platform as paired-end fragments with 2 × 150 bp length at NGX Bio (San Francisco, California, USA). The previously published samples of the DrosRTEC consortium were prepared and sequenced on GAIIX, HiSeq2000 or HiSeq3000 platforms, as described in Bergland *et al*. (2014) and Machado *et al*. (2021). For information on DNA extraction and sequencing methods of the various DGN samples see Lack *et al*. (2016) and others (Pool *et al*. 2012; Langley *et al*. 2012; Grenier *et al*. 2015; Kao *et al*. 2015).

### Mapping pipeline

The joint analysis of genomic data from different sources requires the application of uniform quality criteria and a common bioinformatics pipeline. To accomplish this, we developed a standardized pipeline that performs filtering, quality control and mapping of any given Pool-Seq sample (see supplementary fig. S1). This pipeline performs quality filtering of raw reads, maps reads to a hologenome (see below), performs realignment and filtering around indels, and filters for mapping quality. The output of this pipeline includes quality control metrics, bam files, pileup files, and allele frequency estimates for every site in the genome (gSYNC, see below). Our pipeline is provided as a Docker image and will facilitate the integration of future samples to extend the worldwide *D. melanogaster* SNP dataset presented here.

The mapping pipeline includes the following major steps. Prior to mapping, we removed sequencing adapters and trimmed the 3’ ends of all reads using *cutadapt* (Martin 2011). We enforced a minimum base quality score ≥ 18 (-q flag in *cutadapt*) and assessed the quality of raw and trimmed reads with FASTQC (Andrews 2010). Trimmed reads with minimum length < 75 bp were discarded and only intact read pairs were considered for further analyses. Overlapping paired-end reads were merged using *bbmerge* (v. 35.50; Bushnell *et al*. 2017). Trimmed reads were mapped against a compound reference genome (“hologenome”) consisting of the genomes of *D. melanogaster* (v.6.12) and *D. simulans* (Hu *et al*. 2013) as well as genomes of common commensals and pathogens, including *Saccharomyces cerevisiae* (GCF_000146045.2), *Wolbachia pipientis* (NC_002978.6), *Pseudomonas entomophila* (NC_008027.1), *Commensalibacter intestine* (NZ_AGFR00000000.1), *Acetobacter pomorum* (NZ_AEUP00000000.1), *Gluconobacter morbifer* (NZ_AGQV00000000.1), *Providencia burhodogranariea* (NZ_AKKL00000000.1), *Providencia alcalifaciens* (NZ_AKKM01000049.1), *Providencia rettgeri* (NZ_AJSB00000000.1), *Enterococcus faecalis* (NC_004668.1), *Lactobacillus brevis* (NC_008497.1), and *Lactobacillus plantarum* (NC_004567.2), using *bwa mem* (v. 0.7.15; Li 2013) with default parameters. We retained reads with mapping quality greater than 20 and reads with no secondary alignment using *samtools* (Li *et al*. 2009). PCR duplicate reads were removed using *Picard MarkDuplicates* (v.1.109; http://picard.sourceforge.net). Sequences were re-aligned in the proximity of insertions-deletions (indels) with GATK (v3.4-46; McKenna *et al*. 2010). We identified and removed any reads that mapped to the *D. simulans* genome using a custom python script, following methods outlined previously (Kapun *et al*. 2020; Machado *et al*. 2021; for a more in-depth analysis of *D. simulans* contamination see Wallace *et al*. 2021). Although this method of decontamination by *D. simulans* accurately estimates contamination rate and removes the vast majority of *D. simulans* reads (Machado *et al*. 2021), care should be taken when analyzing samples with higher contamination rates at sites that are shared polymorphisms between the two species.

### Incorporation of the DGN dataset

We incorporated population allele frequency estimates derived from inbred line and haploid embryo sequencing data from populations sampled throughout the world using an *in-silico* pooling approach. These samples have been previously collected and sequenced by several groups (Pool *et al*. 2012; Mackay *et al*. 2012; Langley *et al*. 2012; Grenier *et al*. 2015; Kao *et al*. 2015; Lack *et al*. 2015, 2016) and together form the *Drosophila* Genome Nexus dataset (DGN; Lack *et al*. 2015, 2016). We included 25 DGN populations with ≥ 5 individuals per population, plus the *D. simulans* haplotype w^501^ built as part of the DGN dataset. The DGN populations that we used are primarily from Africa (n=18) but also include populations from Europe (n=2), North America (n=3), Australia (n=1), and Asia (n=1). The complete list of DGN populations, and samples, used in this dataset can be found in supplementary table S1.

To incorporate the DGN populations into the DrosEU and DrosRTEC Pool-Seq datasets, we used the pre-computed FASTA files (“Consensus Sequence Files” from https://www.johnpool.net/genomes.html) and calculated allele frequencies at every site, for each population, using custom *bash* scripts. We calculated allele frequencies for each population by summing reference and alternative allele counts across all individuals using the precomputed haplotype FASTA files. Since estimates of allele frequencies and total allele counts for the DGN samples only consider unambiguous IUPAC codes, heterozygous sites or sites masked as N’s in the original FASTA files were converted to missing data. We used *liftover* (Kuhn *et al*. 2013) to translate genome coordinates to *Drosophila* reference genome release 6 (dos Santos *et al*. 2015) and formatted them to match the gSYNC format (described below). Scripts for reformatting the DGN data can be found in the GitHub repository for this project (https://github.com/DEST-bio/DEST_freeze1).

### SNP calling strategies

We used two complementary approaches to perform SNP calling. The first was PoolSNP (Kapun *et al*. 2020), a heuristic tool which identifies polymorphisms based on the combined evidence from multiple samples. This approach is similar to other common Pool-Seq variant calling tools (Koboldt *et al*. 2009, 2012; Kofler *et al*. 2011a, 2011b). PoolSNP integrates allele counts across multiple independent samples and applies stringent minor allele count and minor allele frequency thresholds for variant detection. PoolSNP is expected to be good at detecting variants present in multiple populations, but is not very sensitive to rare private alleles. The second approach was SNAPE-pooled (Raineri *et al*. 2012), a tool that identifies polymorphic sites based on Bayesian inference for each population independently using pairwise nucleotide diversity estimates as a prior. SNAPE-pooled is expected to be more sensitive to rare private polymorphisms (Rainieri *et al*. 2012, Guirao-Rico and González 2021). The SNP calling step is built using the *snakemake* (Mölder *et al*. 2021) pipeline and the parameters to run the two callers can be found at https://github.com/DEST-bio/DEST_freeze1.

### gSYNC generation and filtering

Our pipeline utilizes a common data format to encode allele counts for each population sample (SYNC; Kofler *et al*. 2011b). A “genome-wide SYNC’’ (gSYNC) file records the number of A,T,C, and G for every site of the reference genome. Because gSYNC files for all populations have the same dimension, they can be quickly combined and passed to a SNP calling tool. They can be filtered and are also relatively small for a given sample (~500 Mb), enabling efficient data sharing and access. The gSYNC file is analogous to the gVCF file format as part of the GATK HaplotypeCaller approach (McKenna *et al*. 2010), but is specifically tailored to Pool-Seq samples.

We generated gSYNC files for both PoolSNP and SNAPE. To generate a PoolSNP gSYNC file, we first converted BAM files to the MPILEUP format with *samtools mpileup* using the -B parameter to suppress recalculations of per-base alignment qualities and filtered for a minimum mapping quality with the parameter -q 25. Next, we converted the MPILEUP file containing mapped and filtered reads to the gSYNC format using custom python scripts. To generate a SNAPE-pooled gSYNC file, we ran the SNAPE-pooled version specific to Pool-Seq data for each sample in MPILEUP format with the following parameters: *θ*=0.005, *D*=0.01, prior=‘informative’, fold=‘unfolded’ and nchr=number of flies (x2 for autosomes and x1 for the X and Y chromosomes) following Guirao-Rico and Gonzalez (2021). We converted the SNAPE-pooled output file to a gSYNC file containing the counts of each allele per position and the posterior probability of polymorphism as defined by SNAPE-pooled using custom python scripts. We only considered positions with a posterior probability ≥ 0.9 as being polymorphic and with a posterior probability ≤ 0.1 as being monomorphic. In all other cases, positions were marked as missing data.

We masked gSYNC files for PoolSNP and SNAPE-pooled using a common set of filters. Sites were filtered from gSYNC files if they had: (1) minimum read depth < 10; (2) maximum read depth > the 95% coverage percentile of a given chromosomal arm and sample; (3) located within repetitive elements as defined by RepeatMasker; (4) within 5 bp distance up- and downstream of indel polymorphisms identified by the GATK IndelRealigner. Filtered sites were converted to missing data in the gSYNC file. The location of masked positions for every sample was recorded as a BED file.

### VCF generation

We generated three versions of the variant files, which differ in their inclusion of the DGN samples and the SNP calling strategy. For PoolSNP variant calling, we generated two variant tables: the first version incorporates all 272 samples of the Pool-Seq (DrosRTEC, DrosEU) and *in-silico* Pool-Seq populations (DGN). The second version only considers the 246 Pool-Seq samples excluding the DGN samples (used for comparison to the SNAPE-pooled version). The third file is based on SNAPE-pooled and contains 246 Pool-Seq samples only.

To generate the PoolSNP versions, we combined the masked PoolSNP-gSYNC files into a two-dimensional matrix, where rows correspond to each position in the reference genome and columns describe chromosome, position and reference allele, followed by allele counts in SYNC format for every sample in the dataset. This combined matrix was then subjected to variant calling using PoolSNP, resulting in a VCF formatted file. We performed SNP calling only for the major chromosomal arms (X, 2L, 2R, 3L, 3R) and the 4th (dot) chromosome. Data for heterochromatic arms of the autosomes, the Y chromosome, and the mitochondrial genome can be extracted from the MPILEUP files provided at https://dest.bio.

We evaluated the choice of two heuristic parameters applied to PoolSNP: global minor allele count (MAC) and global minor allele frequency (MAF). Using all 272 samples, we varied MAF (0.001, 0.01, 0.05) and MAC (5-100) and called SNPs at a randomly selected 10% subset of the genome. Based on SNP annotations with SNPeff (version 4.3; Cingolani *et al*. 2012) we calculated *p*_N_/*p*_S_, which is the ratio of non-synonymous to synonymous polymorphisms, and used this value to tune our choice of MAF and MAC and to identify egregious outlier samples. We found that a global MAC=50 provided qualitatively identical estimates of *p*_N_/*p*_S_ across all populations (Figure 2B) and that the results were insensitive to MAF (results not shown). We therefore used these parameters for genome-wide variant calling (see *Results*: *Identification and quality control of SNP polymorphisms*). We kept a third heuristic parameter, the missing data rate, constant at a minimum of 50%.

To generate the SNAPE-pooled VCF files, we combined the 246 masked SNAPE-pooled gSYNC files into a two-dimensional matrix, as described above, and generated a VCF formatted output based on allele counts for any site found to be polymorphic in one or more populations. We evaluated *p*_N_/*p*_S_ across a range of local minor allele frequency thresholds (Figure 2C) and found that *p*_N_/*p*_S_ is largely insensitive to local MAF, once accounting for some problematic samples (see below).

Final VCF files with annotations from SNPeff (version 4.3; Cingolani *et al*. 2012) were stored in VCF and BCF (Danecek *et al*. 2011) file formats alongside an index file in TABIX format (Li 2011). Besides VCF files, we also stored SNP data in the GDS file format using the *R* package SeqArray (Zheng *et al*. 2017).

### Inversion frequency estimates

We estimated the frequencies of 7 cosmopolitan inversion polymorphisms (*In(2L)t, In(2R)NS, In(3L)P, In(3R)C, In(3R)K, In(3R)Mo, In(3R)Payne*) based on a previously published panel of diagnostic SNP markers that are in tight LD with the corresponding inversions (Kapun *et al*. 2014). As previously described (Kapun *et al*. 2016), we isolated the positions in the VCF file of all marker SNPs and estimated the frequency of each inversion as the mean frequency of inversion-specific alleles at all marker SNPs.

### Population genetic analyses

We estimated allele frequencies for each site across populations as the ratio of the alternate allele count to the total site coverage. We also calculated per-site averages for nucleotide diversity (*π*, Nei 1987), Watterson’s *θ* (Watterson 1975) and Tajima’s *D* (Tajima 1989) across all sites or in non-overlapping windows of 100 kb, 50 kb and 10 kb length. To estimate these summary statistics, we converted masked gSYNC files (with positions filtered for repetitive elements, low and high read depth, and proximity to indels; see *gSYNC generation and filtering*) back to the MPILEUP format using custom-made scripts. mpileup files were processed using npstat v.1 (Ferretti *et al*. 2013) with parameters - maxcov 10000 and -nolowfreq m=0 in order to include all filtered positions for analysis. We only considered sites identified as being polymorphic by PoolSNP or SNAPE-pooled for analysis, using the -snpfile option of npstat. For the DGN populations, chromosome-wide summary statistics were estimated only for samples with less than 50% missing data per chromosome. Due to small sample sizes, Tajima’s *D* was not estimated for 7 African DGN populations that consisted of only 5 haploid embryos. To compare population genetic estimates between the PoolSNP versus SNAPE-pooled datasets, we performed Pearson’s correlations on 226 populations present in both datasets (*see Identification and quality control of SNP polymorphisms*) using the stats package of *R* v.3.6.3. The effects of pool size (number of individuals sampled per population) on genome-wide estimates of *π*, Watterson’s *θ* and Tajima’s *D*_*S*_ estimates were examined for European and North American populations using the PoolSNP dataset and a linear model in *R* v.3.6.3. Finally, for 48 European populations we estimated Pearson’s correlations between *π*, Watterson’s *θ* and Tajima’s *D* as estimated from the PoolSNP dataset versus previous estimates by Kapun *et al*. (2020) using the stats package of *R* v3.6.3.

Next, we examined patterns of between-population differentiation by calculating window-wise estimates of pairwise *F*_ST_, based on the method from Hivert *et al*. (2018) implemented in the computePairwiseFSTmatrix() function of the *R* package poolfstat (v1.1.1). This analysis was performed for the dataset composed of 271 samples (all samples excluding the *D. simulans* reference strain) processed with PoolSNP, focusing on SNPs shared across the whole dataset. Finally, we averaged pairwise *F*_ST_ within and among phylogeographic clusters (Africa [17 samples], North America [76 samples], Eastern Europe [83 samples] and Western Europe [93 samples]; not included due to limited sampling: China and Australia). These *F*_ST_ tracks at windows sizes of 100kb, 50kb and 10kb are available at https://dest.bio (supplementary fig. S2, S3).

To assess population structure in the worldwide dataset, we applied principal components analysis (PCA), population clustering, and population assignment based on a discriminant analysis of principal components (DAPC; Jombart *et al*. 2010) to all 271 PoolSNP-processed samples. For these analyses, we subsampled a set of 100,000 SNPs spaced apart from each other by at least 500 bp. We optimized our models using cross-validation by iteratively dividing the data as 90% for training and 10% for learning. We extracted the first 40 PCs from the PCA and ran Pearson’s correlations between each PC and all loci. We subsequently extracted the top 33,000 SNPs with large and significant correlations to PCs 1-40. We chose the 33,000 number as a compromise between panel size and differentiation power. For example, depending on the number of individuals surveyed, these 33,000 DIMs can discern divergence (*τ*) between two populations with parametric *F*_ST_ of 0.001-0.0001 for sample sizes (n) of 10-1000. These estimates come from the phase change formula: *τ* ≈ *F*_ST_ = 1/(nm)^1/2^ (Patterson *et al*. 2006). Here, the two populations were sampled for n/2 individuals and genotyped at m=33,000 markers. Furthermore, we included SNPs as a function of the percent variance explained by each PC. PCAs, clustering, and assignment-based DAPC analyses were carried out using the *R* packages FactoMiner (v. 2.3), factoextra (v. 1.0.7) and adegenet (v. 2.1.3), respectively.

### Web-based genome browser

Our HTML-based DEST browser (supplementary fig. S2) is built on a JBrowse Docker container (Buels *et al*. 2016), which runs under Apache on a CentOS 7.2 Linux x64 server with 16 Intel Xeon 2.4 GHz processors and 32 GB RAM. It implements a hierarchical data selector that facilitates the visualization and selection of multiple population genetic metrics or statistics for all 271 samples based on the PoolSNP-processed dataset, taking into account sampling location and date. Importantly, our genome browser provides a portal for downloading allelic information and pre-computed population genetics statistics in multiple formats (supplementary fig. S2A, S2C, S3), a usage tutorial (supplementary fig. S2B) and versatile track information (supplementary fig. S2D). Bulk downloads of full variation tracks are available in BigWig format (Kent *et al*. 2010) and Pool-Seq files (in VCF format) are downloadable by population and/or sampling date using custom options from the Tools menu (supplementary fig. S2C). All data, tools, and supporting resources for the DEST dataset, as well as reference tracks downloaded from FlyBase (v.6.12) (dos Santos *et al*. 2015), are freely available at https://dest.bio.

## Results

### Integrating a worldwide collection of *D. melanogaster* population genomics resources

We developed a modular and standardized pipeline for generating allele frequency estimates from pooled resequencing of *D. melanogaster* genomes (supplementary fig. 1). Using this pipeline, we assembled a dataset of allele frequencies from 271 *D. melanogaster* populations sampled around the world (Figure 1A, Supplementary Material online, supplementary table S1). Many of these samples were collected at the same location, at different seasons and over multiple years (Figure 1B). The nature of the genomic data for each population varies as a consequence of biological origin (e.g., inbred lines or Pool-Seq), library preparation method, and sequencing platform.

**Figure 1.**
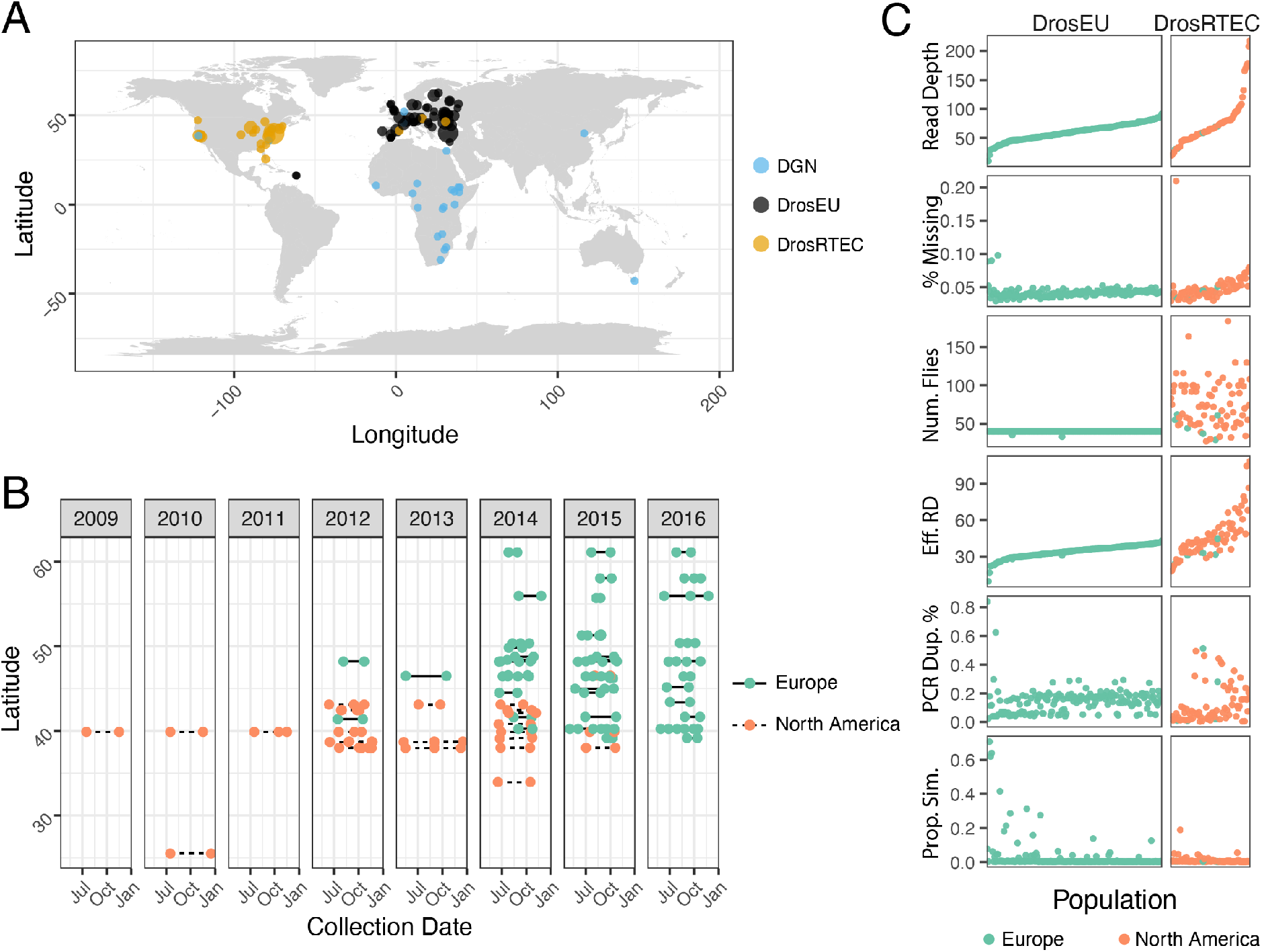
Sampling location, dates, and quality metrics. (A) Map showing the 271 sampling localities forming the DEST dataset. Colors denote the datasets that were combined together. (B) Collection dates for localities sampled more than once. (C) General sample features of the DEST dataset. The x-axis represents the population sample, ordered by the average read depth.

**Figure 2.**
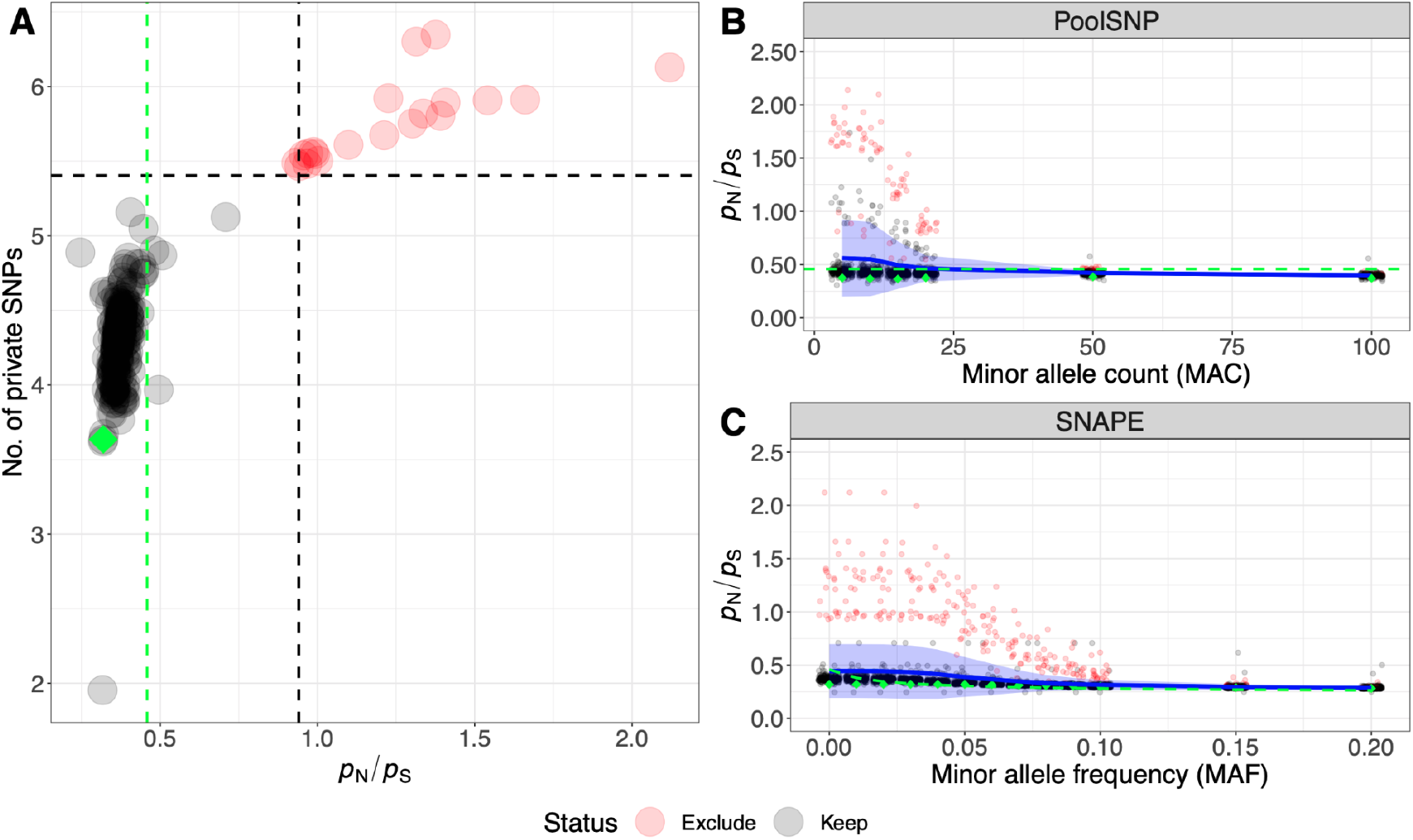
Quality control of SNPs called with SNAPE-pooled and PoolSNP. Panel (A) shows genome-wide *p*_N_/*p*_S_ ratios and the number of private SNPs for all Pool-Seq samples based on SNP calling with SNAPE-pooled. We highlight 20 outlier samples in red, which are characterized by exceptionally high values of both metrics. The dashed black lines indicate the 95% confidence limits (average + 1.96 sd) for both statistics. The vertical green dashed line highlights the empirical estimate of *p*_N_/*p*_S_ calculated from individual sequencing data of the DGRP freeze2 dataset (Mackay *et al*. 2012). The green diamond shows the corresponding value of the DGRP population, which was pool-sequenced as part of the DrosRTEC dataset (NC_ra_03_n; Zhu *et al*. 2012). Panels B and C show the effects of heuristic minor allele count (MAC) and minor allele frequency (MAF) thresholds on *p*_*N*_/*p*_*S*_ ratios in SNP data based on PoolSNP and SNAPE-pooled, respectively. Blue lines in both panels show average genome-wide *p*_N_/*p*_S_ ratios across 271 and 246 populations, respectively. The blue ribbons depict the corresponding standard deviations. The 20 outlier samples, which are characterized in panel A, are highlighted red. In addition, *p*_N_/*p*_S_ ratios of the DGRP Pool-Seq sample (NC_ra_03_n) are shown at different cut-offs as green diamonds and the empirical values from the DGRP freeze2 dataset are indicated as dashed green lines.

To assess whether these features affect basic attributes of the dataset, we calculated six basic quality metrics focusing on the Pool-Seq samples (Figure 1C, Supplementary Material online, supplementary table S2). On average, median read depth across samples is 62x (range: 10-217x). The per-nucleotide missing allele frequency rate was less than 7% for most (95%) of the samples. Excluding populations with high missing data rate (>7%), the proportion of sites with missing data was positively correlated with read depth (*p*=1.2×10^9^, *R*^2^=0.4). The positive correlation between read depth and missing data rate is primarily due to an increased sensitivity to identify indels. The number of flies per sample varied from 33 to 205, with considerable heterogeneity among the DrosRTEC samples (standard deviation [sd]=30), but not among DrosEU samples (sd=0.04). Variation in the number of flies and in sequencing depth is reflected in the effective read depth, an estimate of the number of independent reads after accounting for double binomial sampling that occurs during Pool-Seq (Eff. RD; Kolaczkowski *et al*. 2011; Feder *et al*. 2012; Figure 1C). There was considerable variation in PCR duplicate rate among samples, with notable differences between batches of DrosEU samples (~6% in 2014 vs. 18% in 2015/16; t-test *p*=1.8⨯10^−19^) and DrosRTEC samples (~3% in samples collected as part of Bergland *et al*. 2014 vs. ~14% in samples collected as part of Machado *et al*. 2021; *p*=6.37⨯10^−3^). Curiously, the 2015/2016 DrosEU samples were made with a PCR-free kit, suggesting that the observed PCR duplicates were optical duplicates and not amplification artifacts. Contamination of samples by *D. simulans* varied among populations but was generally absent (<1% *D. simulans* specific reads; supplementary table 1).

### Identification and quality control of SNP polymorphisms

In order to determine appropriate SNP calling and filtering parameters, and to identify potentially problematic population samples, we first calculated the ratio of genome-wide numbers of non-synonymous to synonymous polymorphism (*p*_N_/*p*_S_) for each population sample. Since non-synonymous changes are expected to be under strong purifying selection (Kreitman 1983), we chose this metric because it can reflect the presence of sequencing errors that would disproportionately inflate *p*_N_ relative to *p*_S_. Our primary goal was not to provide novel estimates of *p*_N_/*p*_S_ but rather to ensure that all population samples have estimates that are consistent with estimates generated from independent *Drosophila* datasets (Mackay *et al* 2012).

For the PoolSNP dataset, we varied the global minor allele count (MAC) and global minor allele frequency (MAF) and then calculated *p*_N_/*p*_S_. MAC thresholds <50 resulted in large variances of *p*_N_/*p*_S_ caused by 20 outlier populations characterized by unusually high *p*_N_/*p*_S_ ratios and numbers of private SNPs (Supplementary Material online, supplementary table S3; Figures 2A and 2B) indicating that there may be elevated numbers of sequencing errors in some samples. Some (n=17) of these samples had previously been found to show positive values of Tajima’s *D* across the whole genome (Kapun *et al*. 2020). We observed that, as expected, *p*_N_/*p*_S_ was negatively correlated with MAC (linear regression; *p*<0.001; Figure 2B) and that applying a MAC threshold of 50 reduced the elevated *p*_N_/*p*_S_ ratios of the 20 aforementioned outlier samples to values similar to the rest of the dataset, suggesting that potential sequencing errors had been largely removed. To minimize false positive variant calling, we therefore conservatively chose MAC=50 and MAF=0.001 as threshold parameters for SNP calling with PoolSNP. Using these parameters, we identified 4,381,144 polymorphisms segregating among the 271 *D. melanogaster* samples (Pool-Seq plus DGN), and 4,042,456 polymorphisms segregating among the 246 Pool-Seq samples (excluding DGN), using PoolSNP.

SNAPE-pooled calls variants in each sample separately using a probabilistic approach, in contrast to PoolSNP, which integrates allelic information across all populations for heuristic SNP calling. To quantify the number of putative sequencing errors among low frequency variants we varied the local MAF threshold per sample and calculated *p*_N_/*p*_S_ for each sample in the SNAPE-pooled dataset. Similar to PoolSNP, we found that elevated *p*_N_/*p*_S_ was negatively correlated with a local MAF threshold (linear regression; *p*<0.001; Figure 2C) and that the 20 above-mentioned problematic samples also had a strong effect on the variance and mean of *p*_N_/*p*_S_ ratios. Accordingly, we excluded these 20 samples from further analyses of low-frequency variants and private SNPs and applied a conservative local MAF filter of 5% for the remainder of the SNAPE-pooled analysis to avoid misclassification of sequencing errors as low-frequency variants. Our results identified 8,541,651 polymorphisms segregating among the remaining 226 samples. Below, we discuss the geographic distribution and global frequency of SNPs identified using these two methods in order to provide insight into the marked discrepancy in the number of SNPs that they identify.

### Similarity of SNP polymorphisms detected with PoolSNP and SNAPE-pooled

We calculated three metrics related to the amount of polymorphism discovered by our pipelines: the abundance of polymorphisms segregating in *n* populations across each chromosome (Figure 3A), the difference of discovered polymorphisms between SNAPE-pooled and PoolSNP (defined as the absolute value of PoolSNP minus SNAPE-pooled; Figure 3B), and the amount of polymorphism discovered per minor allele frequency bin (Figure 3C). We evaluated these three metrics across a 2×2 filtering scheme: two MAF filters (0.001, 0.05) and two sample sets (the whole dataset of 246 samples; and the 226 samples that passed the sequencing error filter in SNAPE-pooled; see *Identification and quality control*). Notably, PoolSNP was biased towards identification of common SNPs present in multiple samples, whereas SNAPE-pooled was more sensitive to the identification of polymorphisms that appeared in few populations only (Figure 3B). For example, at a MAF filter of 0.001, SNAPE-pooled discovered more polymorphisms that were shared in less than 25 populations (relative to PoolSNP), and these accounted for ~79% of all polymorphisms discovered by the pipeline. Likewise, at a MAF filter of 0.05, SNAPE-pooled discovered more polymorphisms that were shared in less than 97 populations; these accounted for ~71% of all discovered polymorphisms. SNAPE-pooled identifies fewer polymorphic sites that are shared among a large number of populations than PoolSNP does because SNAPE-pooled does not integrate information across multiple populations. As a consequence, it can fail to identify SNPs that are at low overall frequencies and get called as monomorphic or missing in a subset of populations given the posterior probability thresholds that we employed (see Materials and Methods).

**Figure 3.**
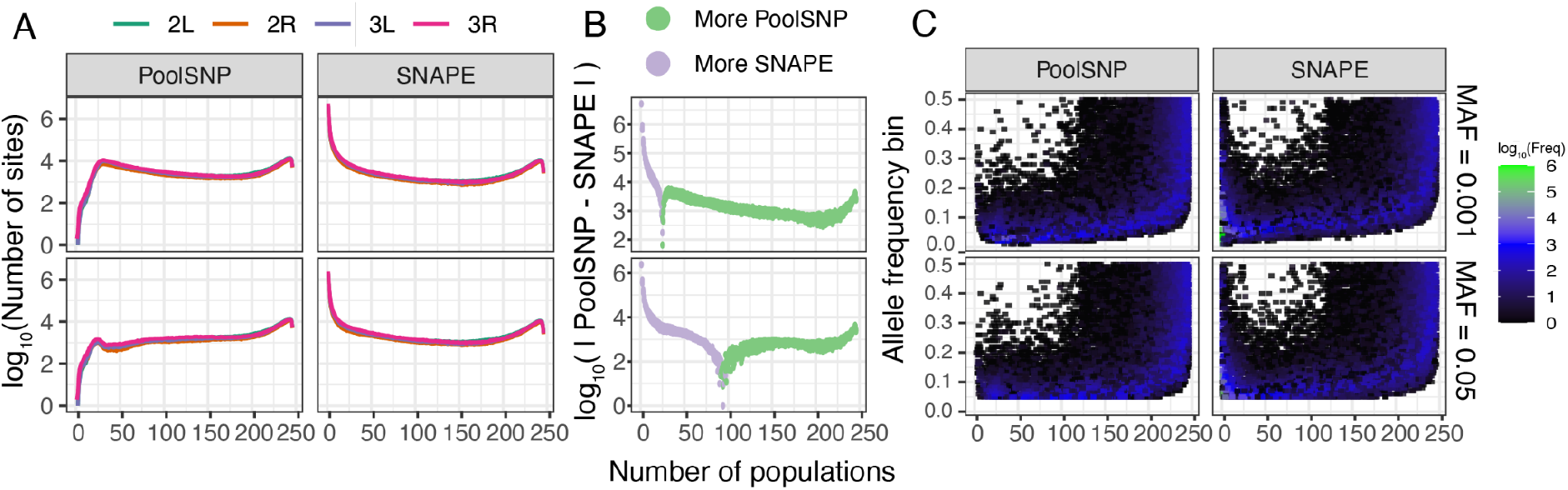
Polymorphism data in the PoolSNP and SNAPE datasets. (A) Number of polymorphic sites discovered across populations. The x-axis shows the number of populations that share a polymorphic site. The y-axis corresponds to the number of polymorphic sites shared by any number of populations, on a log10 scale. The colored lines represent different chromosomes, and are stacked on top of each other. (B) The difference of discovered polymorphisms between SNAPE-pooled and PoolSNP. (C) Number of polymorphic sites as a function of allele frequency and the number of populations in which the polymorphisms are present. The color gradient represents the number of variant alleles from low to high (black to green). The x-axis is the same as in A, and the y-axis is the minor allele frequency. The 2×2 filtering scheme is shown on the right side of the figure.

We also compared allele frequency estimates between the two callers using the aforementioned dataset of 226 populations applying a local MAF filter of 0.05 in the SNAPE-pooled dataset (see Supplementary Material online, supplementary table S2). Among the positions identified as polymorphic by both calling methods, our frequency estimates were identical for the great majority of SNPs (92-99.67%) in all samples analyzed. Between 0.1% and 7.1 % of the polymorphic SNPs differed by less than 5% frequency between the two methods, 0.003 to 2.1% of polymorphic SNPs differed by 5%-10% frequency and only up to 0.3% varied >10% frequency (supplementary table 4). Finally, on average 13.32% of the positions analyzed were called as polymorphic by PoolSNP while there were monomorphic or no data according to SNAPE-pooled, consistent with the use of a hard threshold of the posterior-probability in the SNAPE calling step (Supplementary Material online, supplementary table S4).

### Mutation-class frequencies

We estimated the percentage of mutation classes (e.g., A→C, A→G, A→T, *etc*.) accepted as polymorphisms in both our SNP calling pipelines, and classified these loci as being either “rare” (i.e., allele frequency <5% and shared in less than 50 populations) or “common” (allele frequency >5% and shared in more than 150 populations). For this analysis, we classified the minor allele as the derived allele. Figure 4A shows the percentage of each mutation class for the 226 populations which passed filters in both SNAPE-pooled and PoolSNP. In addition, we overlaid, as a horizontal line, the expected mutation frequencies for rare (blue; Assaf *et al*. 2017) and common (red; Mackay *et al*. 2012) mutations. In general, our SNP discovery pipelines produced mutation-class relative frequencies of rare and common mutations that are consistent with empirical expectations, however, there were some exceptions to this pattern. For example, the frequencies of the C/G rare mutation-class were consistently underestimated by both callers, a phenomenon that might be related to the known GC bias of modern sequencing machines (Benjamini and Speed 2012). The correlation between SNP calling pipelines was high across both common and rare mutation classes, with marginal discrepancies observed for rare variants (Figure 4B).

**Figure 4.**
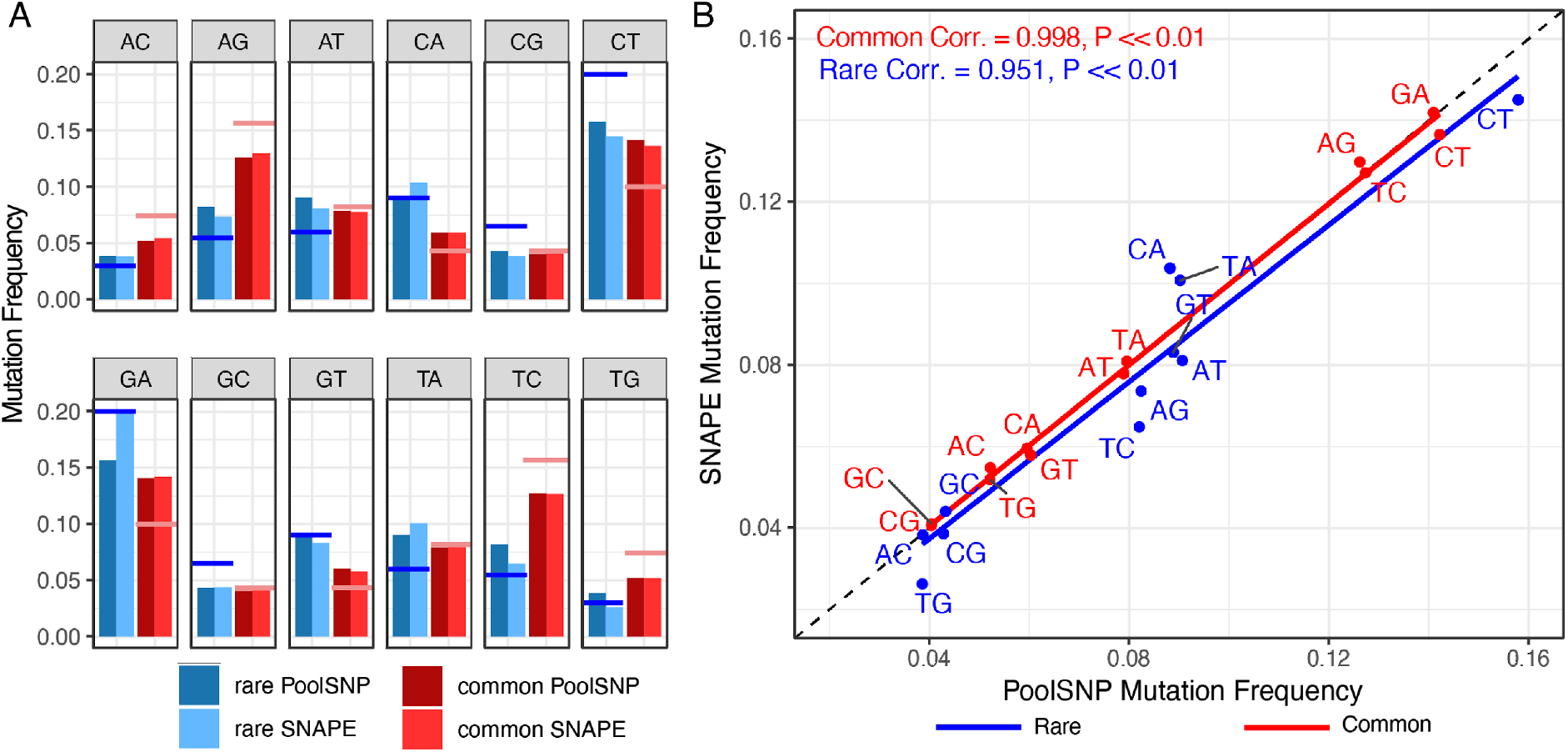
Frequencies of observed nucleotide polymorphism in the DEST dataset (226 populations common to PoolSNP and SNAPE-pooled). (A) Each panel represents a mutation type. The red color indicates common mutations (AF >0.05, and common in more than 150 populations) whereas the blue color indicates rare mutations (AF <0.05, and shared in less than 50 populations). The dark colors correspond to the PoolSNP pipeline and the soft colors correspond to the SNAPE-pooled pipeline. The hovering red and blue horizontal lines represent the estimated mutation rates for common and rare mutations, respectively. (B) Correlation between the observed mutation frequencies seen in SNAPE-pooled and PoolSNP. The one-to-one correspondence line is shown as a black-dashed diagonal. Correlation estimates (Pearson’s correlation) and *p-*values for common and rare mutations are shown.

### Inversion frequencies

Using a set of inversion-specific marker SNPs (Kapun *et al*. 2014), we estimated the frequencies of 7 cosmopolitan inversion polymorphisms (*In(2L)t, In(2R)NS, In(3L)P, In(3R)C, In(3R)K, In(3R)Mo* and *In(3R)Payne*). We found that most of the 271 populations were polymorphic for at least one or more chromosomal inversions (supplementary table 1). While most inversions were either absent or rare (average frequencies: *In(2R)NS* = 5.2% [± 4.7% sd], *In(3L)P* = 3.1% [± 4.3% sd], *In(3R)C* = 2.5% [± 2.3% sd], *In(3R)K* = 1.8% [± 7.4% sd], *In(3R)Mo* = 2.2% [± 3.6% sd] and *In(3R)Payne* = 5.7% [± 7.1% sd]), only *In(2L)t* segregated at substantial frequencies in most populations (average frequency = 18.3% [± 11% sd]). We found that our novel inversion frequency estimates of the DrosEU data from 2014 were highly consistent with previous estimates from Kapun *et al*. (2020) as coefficients of determination (*R*^2^) ranged from 91% to 99%.

### Comparison to previously published datasets

We compared the allele frequency and read depth estimates from the DEST dataset (based on PoolSNP) to previously published estimates by Bergland *et al*. (2014), and Kapun *et al*. (2020), Machado *et al*. (2021). For these datasets we employed two types of correlations, the nominal correlation (i.e., Pearson’s correlation; CO) and the concordance correlation coefficient (CCC; Lin 1989; Liao and Lewis 2000). The CCC determines how much the observed data deviate from the line of perfect concordance (i.e., the 45 degree-line on a square scatter plot).

Estimates of allele frequency were strongly correlated and consistent with previously published data. The strongest correlation of DEST allele frequencies and previously published allele frequencies was observed with the data of Kapun *et al*. (2020) (average CO and CCC >0.99; Figure 5, top row; Supplementary Material online, supplementary fig. S4). Allele frequency correlations with Machado *et al*. (2021) are also generally high (average CO and CCC >0.98; Figure 5, top row; Supplementary Material online, supplementary fig. S5). Allele frequency correlations with the data from Bergland *et al*. (2014) were lower (0.94; Supplementary Material online, supplementary fig. S6), likely reflecting differences in data processing and quality control.

**Figure 5.**
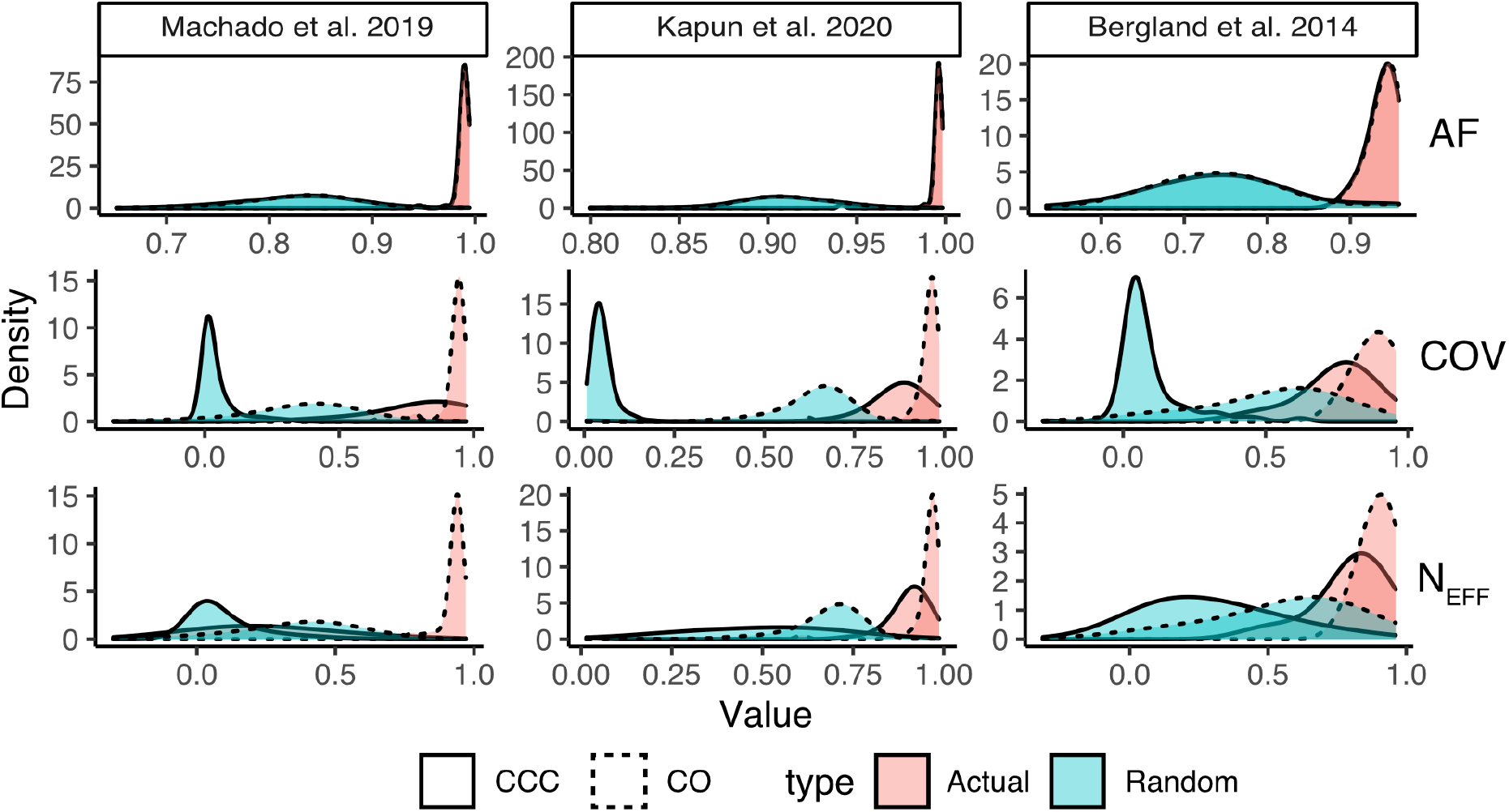
Correlations between DEST dataset and previously published datasets. Correlations between allele frequencies (AF), Nominal Coverage (COV), and Effective Coverage (N_EFF_) between the DEST dataset (using the PoolSNP method) and three previously *Drosophila* datasets: Machado *et al*. (2021), Kapun *et al*. (2020), and Bergland *et al*. (2014). For each dataset, we show the distribution of two types of correlation coefficients: the nominal (Pearson’s) correlation (CO; dashed lines) and the concordant correlation (CCC; solid lines). In addition to the actual correlations between the datasets (red distributions), we show the distributions of correlations estimated with random population pairs (green distributions).

We also examined two aspects of read depth, i.e., nominal coverage and effective coverage. Nominal coverage is the number of reads mapping to a site that has passed quality control. Effective coverage is the approximate number of independent reads, after accounting for double binomial sampling, and is useful for obtaining unbiased estimates of the precision of allele frequency estimates (Kolaczkowski *et al*. 2011; Kofler *et al*. 2011a; Feder *et al*. 2012; Schlötterer *et al*. 2014). Similar to allele frequency estimates, the Pearson correlation coefficients for both coverage and effective coverage were large (0.92, 0.95, 0.90 for Machado *et al*. (2021), Kapun *et al*. (2020), and Bergland *et al*. (2014), respectively; see Supplementary Material online, Figures S7-S12), indicating that sample identity was preserved appropriately. However, the concordance correlation coefficients were substantially lower between the datasets (0.24, 0.88, 0.79, respectively), indicating systematic differences in read depth between the DEST dataset and previously published data. Indeed, read depth estimates were on average ~12%, ~14% and ~20% lower in the DEST dataset as compared to the previously published data in Machado *et al*. (2021), Kapun *et al*. (2020), and Bergland *et al*. (2014) respectively. The lower read depth and effective read depth estimates in the DEST dataset reflect our more stringent quality control and filtering.

### Genetic diversity

We estimated nucleotide diversity (*π*), Watterson’s *θ* and Tajima’s *D* for both the PoolSNP and SNAPE-pooled datasets (Supplementary Material online, supplementary table S5). Results for the African, European and North American population samples are presented in Figure 6 (also see Supplementary Material online, supplementary fig. S13 for estimates by chromosome arm). All estimates were positively correlated between PoolSNP and SNAPE-pooled (*p*<0.001), with Pearson’s correlation coefficients of 0.90, 0.83 and 0.70 for *π*, Watterson’s *θ*, and Tajima’s *D*, respectively. Higher values of genetic diversity were obtained for the SNAPE-pooled dataset, probably due to its higher sensitivity for detecting rare variants (see *Patterns of polymorphism between PoolSNP and SNAPE-pooled*). Pool size had no significant effect on the four summary statistics in European or in North American populations (linear models, all *p*>0.05), suggesting that data from populations with heterogeneous pool sizes can be safely merged for accurate population genomic analysis.

**Figure 6.**
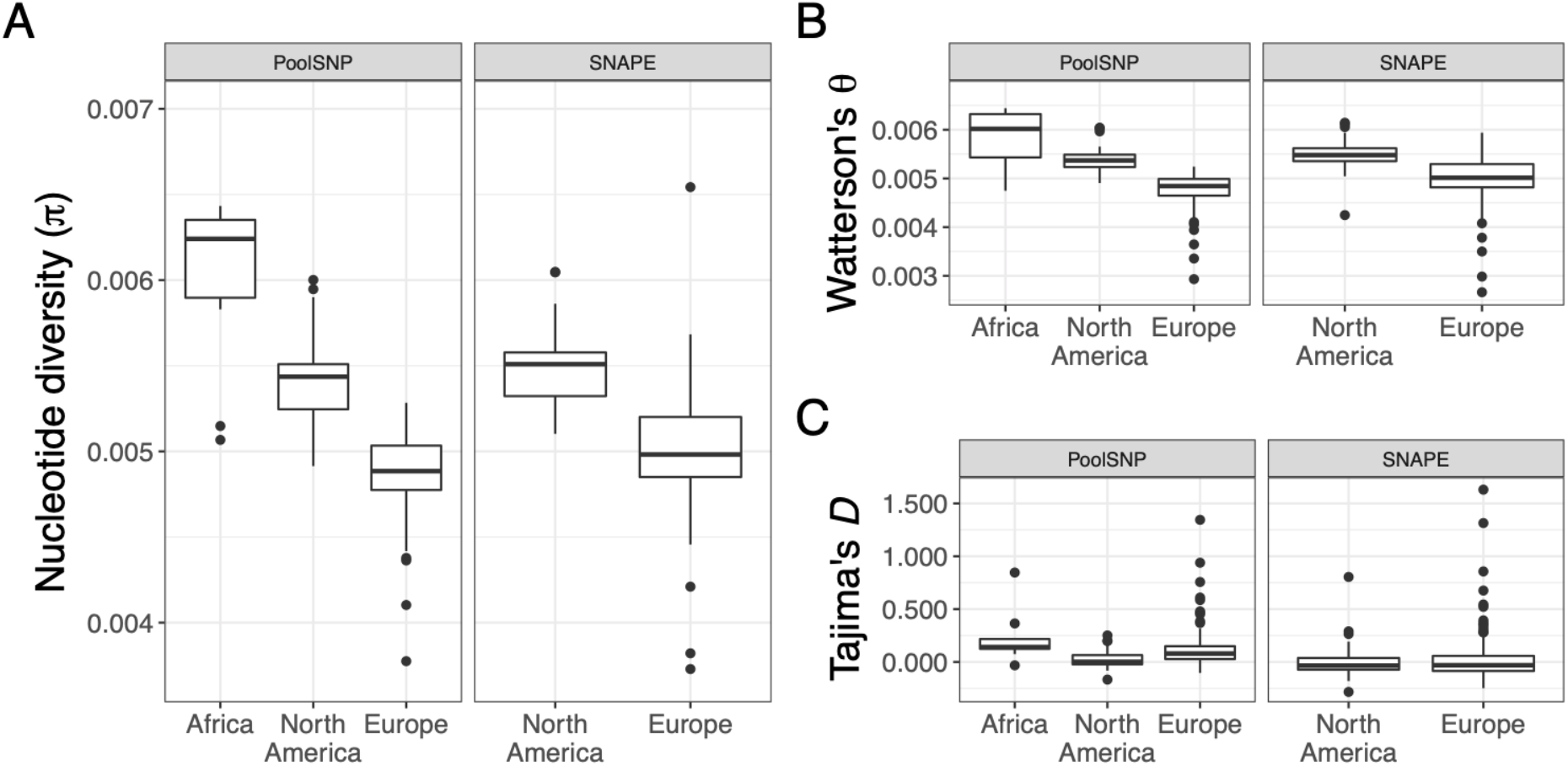
Population genetic estimates for African, European and North American populations. Shown are genome-wide estimates of (A) nucleotide diversity (*π*), (B) Watterson’s *θ* and (C) Tajima’s *D* for African populations using the PoolSNP data set, and for European and North American populations using both the PoolSNP and SNAPE-pooled (SNAPE) datasets. As can be seen from the figure, estimates based on PoolSNP versus SNAPE-pooled (SNAPE) are highly correlated (see main text). Genetic variability is seen to be highest for African populations, followed by North American and then European populations, as previously observed (e.g., see Lack *et al*. 2016; Kapun *et al*. 2020).

The highest levels of genetic diversity were observed for ancestral African populations (mean *π* = 0.0060, mean *θ* = 0.0059); North American populations exhibited higher genetic variability (mean *π* = 0.0054, mean *θ* = 0.0054) than European populations (mean *π* = 0.0049, mean *θ* = 0.0048). These results are consistent with previous observations based on individual genome sequencing (e.g., see Lack *et al*. 2016; Kapun *et al*. 2020). Our observations are also consistent with previous estimates based on pooled data from three North American populations (mean *π* = 0.00577, mean *θ* = 0.00597; Fabian *et al*. 2012) and 48 European populations (mean *π* = 0.0051, mean *θ* = 0.0052; Kapun *et al*. 2020). Estimates of Tajima’s *D* were positive when using PoolSNP, and slightly negative using SNAPE. These results are expected given biases in the detection of rare alleles between these two SNP calling methods. In addition, our estimates for *π*, Watterson’s *θ* and Tajima’s *D* were positively correlated with previous estimates for the 48 European populations analyzed by Kapun *et al*. (2020) (all *p*<0.01). Notably, slightly lower levels of Tajima’s *D* in North America as compared to both Africa and Europe (Figure 6C) may be indicative for admixture (Stajich and Hahn 2005) which has been identified previously along the North American east coast (Caracristi and Schlötterer 2003; Kao *et al*. 2015; Bergland *et al*. 2016).

### Phylogeographic clusters in *D. melanogaster*

We performed PCA on the PoolSNP variants in order to include samples from North America (DrosRTEC), Europe (DrosEU), and Africa (DGN) datasets (excluding all Asian and Oceanian samples). Prior to analysis we filtered the joint datasets to include only high-quality biallelic SNPs. Because LD decays rapidly in *Drosophila* (Comeron *et al*. 2012), we only considered SNPs at least 500 bp away from each other. PCA on the resulting 100,000 SNPs revealed evidence for discrete phylogeographic clusters that correspond to geographic regions (Supplementary Material online, supplementary fig. S14B). PC1 (24% variance explained [VE]) partitions samples between Africa and the other continents (Figure 7A). PC2 (9% VE) separates European from North American populations, and both PC2 and PC3 (4% VE) divide Europe into two population clusters (Figure 7B). As expected, North American samples are intermediate to European and African samples, presumably due to recent secondary contact (Kao *et al*. 2015; Pool 2015; Bergland *et al*. 2016). Notably, these spatial relationships become evident when PCA projections from each sample are plotted onto a world map (Figure 7C). Interestingly, the emergent clusters in Europe are not strictly defined by geography. For example, the western cluster (diamonds in Figure 7D) includes Western Europe as well as Finland, Turkey, Cyprus, and Egypt. The eastern cluster, on the other hand, consists of several populations collected in previous Soviet republics as well as Poland, Hungary, Serbia and Austria, raising the possibility that recent geo-political division in Europe could have affected migration and population structure. Whether this result arises as a relic of recent geopolitical history within Europe, more ancient migration and colonization (e.g., following post-glacial range expansion, Kapun *et al*. 2020), local adaptation, or sampling strategy (Novembre and Stephens 2008; cf. Kapun *et al*. 2020) remains unknown. Future targeted sampling is needed to resolve these alternative explanations.

**Figure 7.**
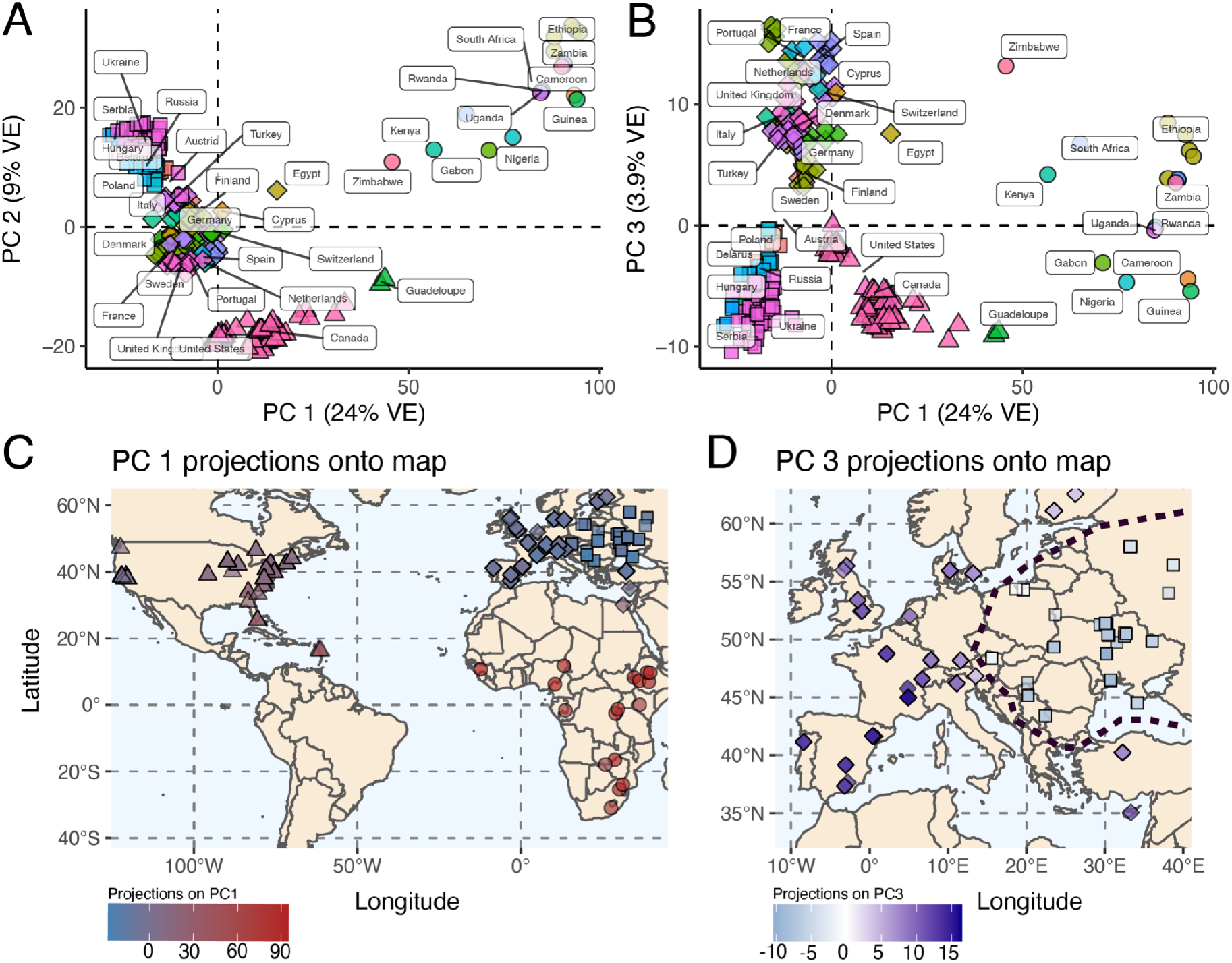
Demographic signatures of the DrosEU, DrosRTEC, and DGN data (using the PoolSNP pipeline). (A) PCA dimensions 1 and 2. The mean centroid of a country’s assignment is labeled. (B) PCA dimensions 1 and 3. (C) Projections of PC1 onto a World map. PC1 projections define the existence of continental level clusters of population structure (indicated by the shapes circles: Africa; triangles: North America; diamonds and squares: Europe). (D) Projections of PC3 onto Europe. These projections show the existence of a demographic divide within Europe: the diamond shapes indicate a western cluster, whereas the squares represent an eastern cluster. For panels C and D, the intensity of the color is proportional to the PC projection. The black dashed line shows the two-cluster divide.

A unique feature of this dataset is that it contains a mixture of Pool-Seq and inbred (or haploid) genome data. For some geographic regions, the DEST dataset contains both data types. Inbred and Pool-Seq samples from nearby geographic regions clustered in the same regions of PC space (Supplementary Material online, supplementary fig. S15). Excluding the DGN-derived African samples, no PC was significantly correlated with data type (PC1 *p* = 0.352, PC2 *p* = 0.223, PC3 *p* = 0.998).

### Geographic proximity analysis

The geographic distribution of our samples allows leveraging basic principles of phylogeography and population genetics to assess the biological significance of rare SNPs (Wright 1943; Battey *et al*. 2020). Accordingly, we expect to observe young neutral alleles at low frequencies among geographically close populations, reflecting isolation by distance. We tested this hypothesis by estimating the average geographic distance among pairs of populations that share SNPs only occurring in these two populations (doubletons), among three populations that share tripletons, and so forth. Without imposing a MAF filter, both SNAPE-pooled and PoolSNP pipelines produced patterns concordant with the expectation. Populations in close proximity were more likely to share rare mutations relative to random chance pairings (Figure 8A). Notably, SNPs identified in less than 25 populations tend to be geographically closer in PoolSNP, relative to SNAPE-pooled. The primary source of this discrepancy between callers occurs when evaluating SNPs shared by just 2 populations (Figure 8B). In the case of PoolSNP, only 0.0006% of all SNPs are private to just 2 populations and the mean geographical distance is 702 Km. In the case of SNAPE-pooled, 9.3% of all SNPs are private to 2 populations and the mean distance is ~2000 Km. Aside from the case of n=2, the difference in proximity estimates between the callers is minimal. These findings suggest that some of the SNAPE-pooled SNPs which only segregate in two populations or less might be false positives. To further evaluate these geographical patterns, we estimated the probability that any given population pair belongs to a particular phylogeographic cluster (Supplementary Material online, supplementary fig. S16) as a function of their shared variants. Our results indicate that rare variants, private to geographically proximate populations, are strong predictors of phylogeographic provenance (see Figure 8C).

**Figure 8.**
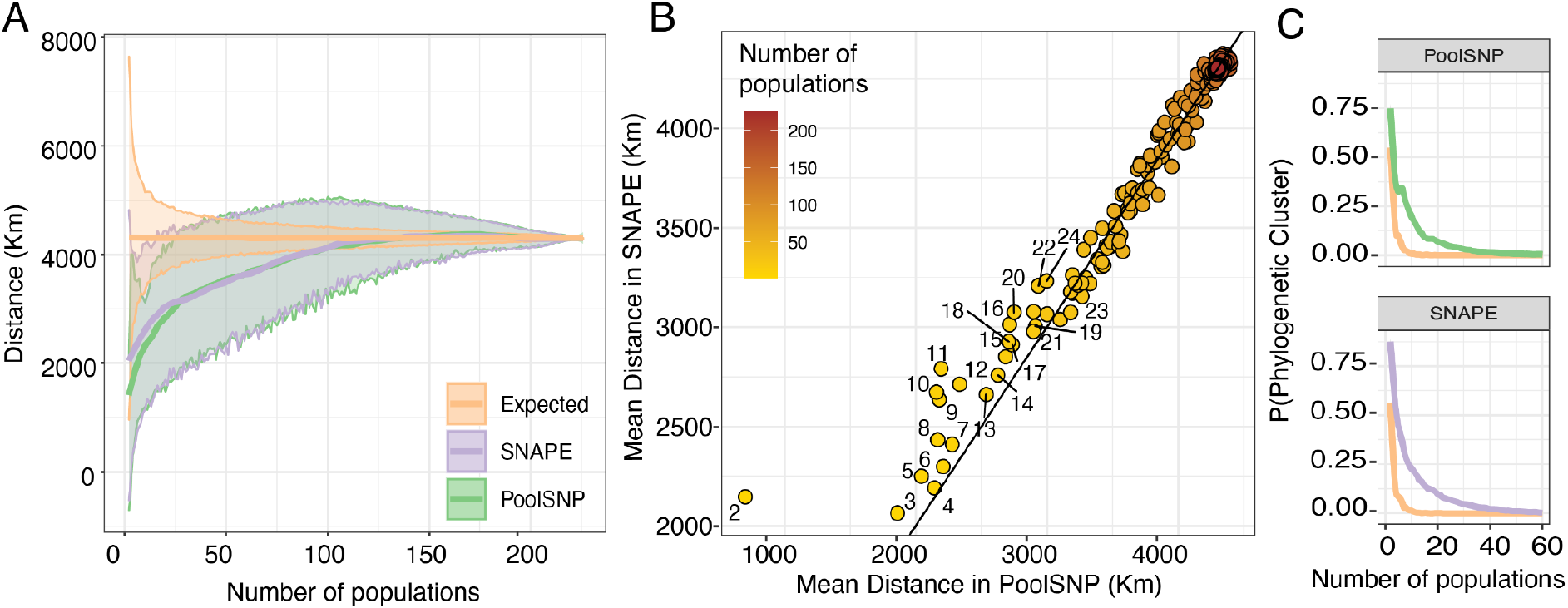
Geographic Proximity Analysis. (A) Average (local regression; LOESS) geographic distance between populations that share a polymorphism at any given site for PoolSNP and SNAPE-pooled. The x-axis represents the number of populations considered; the y-axis is the mean geographic distance among samples. The yellow line represents the random expectation calculated as random pairings of the data. The band around the lines is the standard deviation of the estimator. (B) Correlation graph showing the different mean distance estimate for both callers as a function of the number of populations (the groups from n=2 to n=25 are labeled in the graph). A 1-to-1 line is also shown. (C) Probability that all populations containing a polymorphic site come from the same phylogeographic cluster (as defined by principal component space, Figure 7 and supplementary fig. 14). The y-axis is the probability of “x” populations belonging to the same phylogeographic cluster. The axis only shows up to 60 populations since, after 40 populations, the probabilities approach 0. The colors are consistent across panels.

### Geographically-informative markers

An inherent strength of our broad biogeographic sampling is the potential to generate a panel of core demography SNPs to investigate the provenance of current and future samples. We created a panel of geographically-informative markers (GIMs) by conducting a discriminant analysis of principal components (DAPC) to discover which loci drive the phylogeographic signal in the dataset. We trained two separate DAPC models: the first utilized the four phylogeographic clusters identified by principal components (PCs; Figure 6AB, Supplementary Material online, supplementary fig. S16, supplementary table S1); the second utilized the geographic localities where the samples were collected (i.e., countries in Europe and the US states). This optimization indicated that the information contained in the first 40 PCs maximizes the probability of successful assignment (Figure 9A). This resulted in the inclusion of 30,000 GIMs, most of which were strongly associated with PCs 1-3 (Figure 9B inset). Moreover, the correlations were larger among the first 3 PCs and decayed monotonically for the additional PCs (Figure 9B). Lastly, our GIMs were uniformly distributed across the fly genome (Figure 9C).

**Figure 9.**
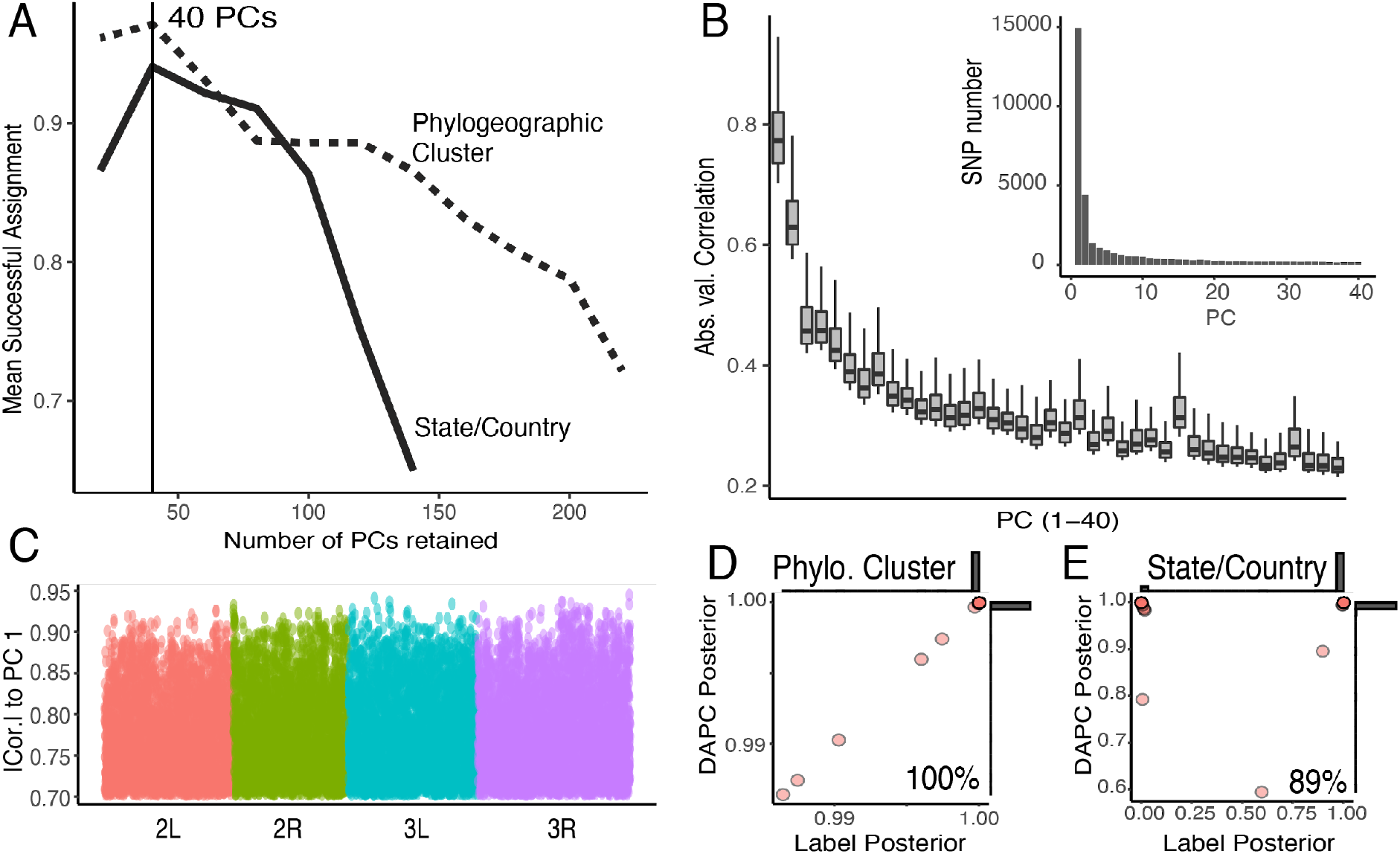
Geographically-informative markers. (A) Number of retained PCs which maximize the DAPC model’s capacity to assign group membership. Model trained on the phylogeographic clusters (dashed lines) or the country/state labels (solid line). (B) Absolute correlation for the 33,000 individual SNPs with highest weights onto the first 40 components of the PCA. Inset: Number of SNPs per PC. (C) Location of the 33,000 most informative demographic SNPs across the chromosomes. (D) LOOCV of the DAPC model trained on the phylogeographic clusters. (E) LOOCV of the DAPC model trained on the phylogeographic state/country labels. For panels D and E, the y-axis shows the highest posterior produced by the prediction model and the x-axis is the posterior assigned to the actual label classification of the sample. Also, for D and E, marginal histograms are shown.

We assessed the accuracy of our GIM panel using a leave-one-out cross-validation approach (LOOCV). We trained the DAPC model using all but one sample and then classified the excluded sample. We performed LOOCV separately for the phylogeographic cluster groups, as well as for the state/country labels. The phylogeographic model used all DrosRTEC, DrosEU, and DGN samples (excluding Asia and Oceania with too few individuals per sample); the state/country model used only samples for which each label had at least 3 or more samples. Our results showed that the model is 100% accurate in terms of resolving samples at the phylogeographic cluster level (Figure 9D) and 89% at the state/country level (Figure 9E). We anticipate that this set of DIMs will be useful to validate the geographic origin of samples in future sequencing efforts (i.e., identify sample swaps; Nunez *et al*. 2021) and to study patterns of migration. We note that although *Drosophila* populations evolve over short time-scales in temperate orchards, samples collected over multiple years were predicted with 89% accuracy in our LOOCV analysis, suggesting that these markers will be valuable for future samples. We provide a tutorial on the usage of the GIMs in Supplemental Methods.

## Discussion

Here we have presented a new, modular and unified bioinformatics pipeline for processing, integrating and analyzing SNP variants segregating in population samples of *D. melanogaster*. We have used this pipeline to assemble the largest worldwide data repository of genome-wide SNPs in *D. melanogaster* to date, based both on previously published data (DGN: Africa; Lack *et al*. 2015, 2016) as well as on new data collected by our two collaborating consortia (DrosRTEC: mostly North America; Machado *et al*. 2021; DrosEU: mostly Europe; Kapun *et al*. 2020). We assembled this dataset using two SNP calling strategies that differ in their ability to identify rare polymorphisms, thereby enabling future work studying the evolutionary history of this species. We are dubbing this data repository and the supporting bioinformatics tools *Drosophila Evolution over Space and Time* (DEST).

The DEST data repository was built using two different SNP calling pipelines, SNAPE-pooled (Raineri *et al*. 2012) and PoolSNP (Kapun *et al*. 2020). These two approaches differ fundamentally in their approach to SNP identification. SNAPE-pooled treats each Pool-Seq sample separately and calculates the posterior probability that a site is polymorphic based on the read depth, alternate allele count, and a prior estimate of nucleotide diversity; this approach was designed to identify rare polymorphisms and has been validated using both simulation and empirical approaches (Guirao-Rico and Gonzalez 2021). Here, we also provide evidence that rare and private SNPs identified by SNAPE-pooled are enriched for true positives (Figure 8) after applying rigorous filtering and excluding 20 population samples likely affected by problems during library preparation which may have resulted in elevated error rates.

The dataset based on SNAPE-pooled could therefore be useful for studies that rely on rare SNPs, such as those investigating recent demographic events (Keinan and Clark 2012). SNAPE-pooled has several limitations though. First, it is only capable of handling Pool-Seq data. Second, because of the hard-filtering that we are imposing with our posterior probability cutoff, some true SNPs are being called as missing data (see Materials and Methods). This problem is apparent when comparing the number of polymorphisms identified by SNAPE-pooled and PoolSNP (Figure 3). In addition, studies that rely on the SNAPE-pooled dataset should exclude the 20 samples we flagged here (Figure 2A, supplementary table 1).

PoolSNP, on the other hand, is useful for analysis of common variants and allows studying aspects of population structure and local adaptation based on shared polymorphism. Such analyses could include the inference of migration out of Africa Kapopoulou *et al*. 2020), admixture (Bergland *et al*. 2016), and back-migration to Africa (Pool and Aquadro 2006). PoolSNP is an extension of the approach developed elsewhere (Kofler *et al*. 2011a,b; Kapun *et al*. 2020). PoolSNP necessarily has a limited capacity to identify rare and private SNPs because it imposes global minor allele count and allele frequency filters. As a consequence, the more populations that are used for SNP calling by PoolSNP, the less likely PoolSNP is to identify private polymorphisms. Because PoolSNP filters out rare and private polymorphisms, it is less sensitive to sequencing or library preparation errors. Notably, the 20 flagged populations do not have elevated *p*_N_/*p*_S_ with MAC > 50. Additionally, Kapun *et al*. (2020) demonstrated that these problematic samples did not affect population genetic inference based on common SNPs. The problematic samples derived from the DrosRTEC studies likely do not have a major impact on their results either as both Bergland *et al*. (2014) and Machado *et al*. (2021) imposed stringent minor allele frequency filters.

PoolSNP has the advantage that it can incorporate in-silico pooled datasets wherein haplotype or genotype information are collapsed into allele frequencies (see Materials and Methods). We took this approach by incorporating the *Drosophila* Genome Nexus dataset (DGN; Lack *et al*. 2016), a dataset that amalgamates whole-genome sequencing of inbred line data and haploid embryos from samples collected around the world. Although the DGN data was originally generated by multiple labs and run through a different mapping pipeline than what we used for the Pool-Seq data, these samples appear to cluster tightly with geographically close Pool-Seq samples (supplementary fig. S15, and discussed in the Results). Thus, there does not appear to be significant bias when combining these datasets, at least when integrating information across the genome. Nonetheless, some care should be taken when interpreting allele frequency differences based on datasets generated by different means. However, any real-time monitoring activity will likely suffer from the rapidly changing landscape of sequencing technologies.

One of the biggest challenges in the present “omics” era is the rapidly growing number of complex large-scale datasets which require technically elaborate bioinformatics know-how to become accessible and utilizable. This hurdle often prohibits the exploitation of already available genomics datasets by scientists without a strong bioinformatics or computational background. To remedy this situation for the *Drosophila* evolution community, our bioinformatics pipeline is provided as a Docker image (to standardize across software versions, as well as make the pipeline independent of specific operating systems) and a new genome browser makes our SNP dataset available through an easy-to-use web interface (see supplementary fig. S2, S3; available at https://dest.bio).

The DEST data repository and platform will enable the population genomics community to address a variety of longstanding, fundamental questions in ecological and evolutionary genetics. The current dataset might for instance be valuable for providing a more accurate picture of the demographic history of *D. melanogaster* populations, in particular in Europe and North America, and with respect to multiple bouts of out-of-Africa migration and recent patterns of admixture. Such analyses can be strongly affected by chromosomal inversions that are known to impact LD and haplotype variation (Kapun and Flatt 2019; Durmaz *et al*. 2020). We have therefore provided frequency estimates for the seven most common cosmopolitan inversions (*In(2L)t, In(2R)NS, In(3L)P, In(3R)C, In(3R)K, In(3R)Mo* and *In(3R)Payne*; Lemeunier and Aulard 1992), which allows accounting for the effects of inversions in population genetic inference (e.g., Kapopoulou *et al*. 2020).

The DEST dataset will likewise be useful for an improved understanding of the genomic signatures underlying both global and local adaptation, including a more fine-grained view of selective sweeps, their evolutionary origin and distribution (e.g., see Glinka *et al*. 2003; Beisswanger *et al*. 2006; Ometto 2010; Stephan 2016; Kapun *et al*. 2020). In terms of local adaptation, the broad spatial sampling across latitudinal and longitudinal gradients on the North American and European continents, encompassing a broad range of climate zones and areas of varying degrees of seasonality, will allow examining the parallel nature of local (clinal) adaptation in response to similar environmental factors in greater depth than possible before (e.g., Turner *et al*. 2008; Kolaczkowski *et al*. 2011; Fabian *et al*. 2012; Reinhardt *et al*. 2014; Bergland *et al*. 2014, 2016; Kapun *et al*. 2016, 2020; Bogaerts-Márquez *et al*. 2020; Waldvogel *et al*. 2020; Machado *et al*. 2021).

Another major opportunity provided by the DEST dataset lies in studying the temporal dynamics of evolutionary change. Sampling at dozens of localities across the growing season and over multiple years will help to advance our understanding of the short-term population and evolutionary dynamics of flies living in diverse environments, thereby providing novel insights into the nature of temporally varying selection (Bergland *et al*. 2014; Wittmann *et al*. 2017; Machado *et al*. 2021) and evolutionary responses to climate change (e.g., Umina 2005; Rodríguez-Trelles *et al*. 2013; Waldvogel *et al*. 2020).

Moreover, by integrating these worldwide estimates of allele frequencies, those from lab- and field-based ‘evolve and resequence’ experiments (E&R; Turner *et al*. 2011; reviewed in Kofler and Schlötterer 2014; Schlötterer *et al*. 2014; Flatt 2020) and those from mesocosm experiments (e.g., Rudman *et al*. 2019; Erickson *et al*. 2020), we might be able to gain deeper insights into the genetic basis and evolutionary history of variation in fitness components (e.g., Flatt 2020).

The real value of the DEST dataset lies in the future: its long-term utility will grow as natural and experimental populations are continually being sampled, resequenced and added to the repository by the community of *Drosophila* evolutionary geneticists. The pipeline that we have established will make future updates to the data repository straightforward. Furthermore, since it is not easily feasible for any single research group to sample flies densely through time and across a broad geographic range, the growing value of the DEST dataset will depend upon the synergistic collaboration among research groups across the globe, as exemplified by the DrosRTEC and DrosEU consortia. Importantly, in an era of rapidly decreasing sequencing costs, comprehensive population genomic analyses are no longer limited by genetic marker density but by the availability of biological samples from standardized, collaborative long-term collection efforts through space and time (e.g., Kapun *et al*. 2020; Machado *et al*. 2021). In this vein, the collaborative framework presented here might allow us, as a global community, to fill some important gaps in the current data repository: for example, many areas of the world (notably Asia and South America) remain largely uncharted territory in *Drosophila* population genomics, and the addition of phased sequencing data (e.g., providing information on haplotypes, LD, linked selection) will be crucially important for future analyses of demography, selection and their interplay.

We are convinced that the DEST platform will become a valuable and widely-used resource for scientists interested in *Drosophila* evolution and genetics, and we actively encourage the community to join the collaborative effort we are seeking to build.

## Data availability

All scripts to make figures and perform analyses associated with this manuscript are available here: https://github.com/DEST-bio/data-paper. All scripts to build the dataset, including the mapping pipeline, SNP calling scripts, and meta-data are available here: https://github.com/DEST-bio/DEST_freeze1. All output from the DEST pipeline, including intermediate output files, metadata, etc. can be found here: https://dest.bio. Datafiles available via the website can also be downloaded through the command-line interface. The genome browser associated with the DEST dataset can be found here: https://dest-bio.uab.cat. The dockerized mapping pipeline can be found at https://hub.docker.com/orgs/destbio/repositories.

## Acknowledgements

We thank four reviewers and the handling editor for helpful comments on a previous version of our manuscript. We are grateful to the members of the DrosEU and DrosRTEC consortia for their long-standing support, collaboration and for discussion. DrosEU is funded by a Special Topic Networks (STN) grant from the European Society for Evolutionary Biology (ESEB). MK (M. Kapun) was supported by the Austrian Science Foundation (grant no. FWF P32275); JG by the European Research Council (ERC) under the European Union’s Horizon 2020 research and innovation programme (H2020-ERC-2014-CoG-647900) and by the Spanish Ministry of Science and Innovation (BFU-2011-24397); TF by the Swiss National Science Foundation (SNSF grants PP00P3_133641, PP00P3_165836, and 31003A_182262) and a Mercator Fellowship from the German Research Foundation (DFG), held as a EvoPAD Visiting Professor at the Institute for Evolution and Biodiversity, University of Münster; AOB by the National Institutes of Health (R35 GM119686); MK (M. Kankare) by Academy of Finland grant 322980; VL by Danish Natural Science Research Council (FNU) grant 4002-00113B; FS Deutsche Forschungsgemeinschaft (DFG) grant STA1154/4-1, Project 408908608; JP by the Deutsche Forschungsgemeinschaft Projects 274388701 and 347368302; AU by FPI fellowship (BES-2012-052999); ET Israel Science Foundation (ISF) grant 1737/17; MSV, MSR and MJ by a grant from the Ministry of Education, Science and Technological Development of the Republic of Serbia (451-03-68/2020-14/200178); AP, KE and MT by a grant from the Ministry of Education, Science and Technological Development of the Republic of Serbia (451-03-68/2020-14/200007); and TM NSERC grant RGPIN-2018-05551. The authors acknowledge Research Computing at The University of Virginia for providing computational resources and technical support that have contributed to the results reported within this publication (https://rc.virginia.edu).

## Author contributions

<u>Martin Kapun</u>: Conceptualization, Data curation, Formal Analysis, Funding acquisition, Investigation, Methodology, Project Administration, Resources, Software, Supervision, Visualization, Writing - original draft, Writing - review & editing. <u>Joaquin Nunez</u>: Formal Analysis, Methodology, Software, Visualization, Writing - original draft, Writing - review & editing. <u>María Bogaerts-Márquez</u>: Formal Analysis, Methodology, Software, Visualization, Writing - original draft, Writing - review & editing. <u>Jesús Murga-Moreno</u>: Formal Analysis, Methodology, Software, Visualization, Writing - original draft, Writing - review & editing. <u>Margot Paris</u>: Formal Analysis, Methodology, Software, Visualization, Writing - original draft, Writing - review & editing. <u>Joseph Outten</u>: Software, Writing - review & editing. <u>Marta Coronado-Zamora</u>: Formal Analysis, Methodology, Software, Visualization, Writing - original draft, Writing - review & editing. <u>Aleksandra Patenkovic</u>: Resources. <u>Amanda Glaser-Schmitt</u>: Resources. <u>Anna Ullastres</u>: Resources. <u>Antonio J. Buendía-Ruíz</u>: Resources. <u>Banu S. Onder</u>: Resources. <u>Brian P Lazzaro</u>: Resources, Writing - review & editing. <u>Catherine Montchamp-Moreau</u>: Resources. <u>Christopher W. Wheat</u>: Resources, Writing - review & editing. <u>Cristina P. Vieira</u>: Resources, Writing - review & editing. <u>Daniel K. Fabian</u>: Resources. <u>Darren J. Obbard</u>: Resources. <u>Dmitry V. Mukha</u>: Resources. <u>Dorcas J. Orengo</u>: Resources, Writing - review & editing. <u>Elena Pasyukova</u>: Resources. <u>Eliza Argyridou</u>: Resources. <u>Emily L. Behrman</u>: Resources, Writing - review & editing. <u>Eran Tauber</u>: Resources. <u>Eva Puerma</u>: Resources, Writing - review & editing. <u>Fabian Staubach</u>: Resources, Writing - review & editing. <u>Francisco D Gallardo-Jiménez</u>: Resources. <u>Iryna Kozeretska</u>: Resources. J. <u>Roberto Torres</u>: Resources. <u>Jessica K. Abbott</u>: Resources. <u>John Parsch:</u> Funding acquisition, Resources, Writing - review & editing. <u>Jorge Vieira</u>: Resources, Writing - review & editing. <u>M. Josefa Gómez-Julián</u>: Resources. <u>Katarina Eric</u>: Resources. <u>Kelly A. Dyer</u>: Resources. <u>Lain Guio</u>: Resources. <u>Lino Ometto</u>: Writing - review & editing. <u>M. Luisa Espinosa-Jimenez</u>: Resources. <u>Maaria Kankare</u>: Resources, Writing - review & editing. <u>Mads F. Schou</u>: Resources, Writing - review & editing. <u>Maria P. García Guerreiro</u>: Resources, Writing - review & editing. <u>Marija Savic Veselinovic</u>: Resources. <u>Marija Tanaskovic</u>: Resources. <u>Marina Stamenkovic-Radak</u>: Funding acquisition, Resources. <u>Mihailo Jelic</u>: Resources. <u>Miriam Merenciano</u>: Resources. <u>Oleksandr M. Maistrenko</u>: Writing - review & editing. <u>Omar Rota-Stabelli</u>: Resources. <u>Sara Guirao-Rico</u>: Resources, Writing - review & editing. <u>Sònia Casillas</u>: Resources, Writing - review & editing. <u>Sonja Grath</u>: Resources. <u>Stephen W. Schaeffer</u>: Resources. <u>Subhash Rajpurohit</u>: Resources. <u>Svitlana V. Serga</u>: Resources. <u>Thomas J.S. Merritt</u>: Resources. Vivien Horváth: Resources. <u>Vladimir E. Alatortsev</u>: Resources. <u>Volker Loeschcke</u>: Resources. <u>Yun Wang</u>: Resources. <u>Heather E. Machado</u>: Resources. <u>Antonio Barbadilla</u>: Software, Writing - review & editing. <u>Dmitri Petrov</u>: Conceptualization, Funding acquisition, Project Administration, Resources, Writing - review & editing. <u>Paul Schmidt</u>: Conceptualization, Funding acquisition, Project Administration, Resources, Writing - review & editing. <u>Josefa Gonzalez</u>: Conceptualization, Funding acquisition, Project Administration, Resources, Supervision, Writing - original draft, Writing - review & editing. <u>Thomas Flatt</u>: Conceptualization, Funding acquisition, Project Administration, Resources, Supervision, Writing - original draft, Writing - review & editing. <u>Alan Bergland</u>: Conceptualization, Data curation, Formal Analysis, Funding acquisition, Investigation, Methodology, Project Administration, Resources, Software, Supervision, Visualization, Writing - original draft, Writing - review & editing

## Competing interests

The authors declare no competing interests.

## Notes

https://dest.bio

### Competing Interest Statement

The authors have declared no competing interest.

## References

Adams, M. D., 2000 The Genome Sequence of Drosophila melanogaster. Science 287: 2185–2195.

Andrews, S., 2010 FastQC: A Quality Control Tool for High Throughput Sequence Data.

Arguello, J. R., S. Laurent, and A. G. Clark, 2019 Demographic History of the Human Commensal Drosophila melanogaster. Genome Biology and Evolution 11: 844–854.

Assaf, Z. J., S. Tilk, J. Park, M. L. Siegal, and D. A. Petrov, 2017 Deep sequencing of natural and experimental populations of Drosophila melanogaster reveals biases in the spectrum of new mutations. Genome Research 27: 1988–2000.

Bastide, H., A. Betancourt, V. Nolte, R. Tobler, P. Stöbe et al., 2013 A Genome-Wide, Fine-Scale Map of Natural Pigmentation Variation in Drosophila melanogaster. PLoS Genet 9: e1003534.

Battey, C. J., P. L. Ralph, and A. D. Kern, 2020 Space is the Place: Effects of Continuous Spatial Structure on Analysis of Population Genetic Data. Genetics 215: 193–214.

Behrman, E. L., V. M. Howick, M. Kapun, F. Staubach, A. O. Bergland et al., 2018 Rapid seasonal evolution in innate immunity of wild Drosophila melanogaster. Proceedings of the Royal Society B: Biological Sciences 285: 20172599.

Beisswanger, S., W. Stephan, and D. Lorenzo, 2006 Evidence for a Selective Sweep in the wapl Region of Drosophila melanogaster. Genetics 172: 265–274.

Benjamini, Y., and T. P. Speed, 2012 Summarizing and correcting the GC content bias in high-throughput sequencing. Nucleic Acids Research 40: e72.

Bergland, A. O., E. L. Behrman, K. R. O’Brien, P. S. Schmidt, and D. A. Petrov, 2014 Genomic Evidence of Rapid and Stable Adaptive Oscillations over Seasonal Time Scales in Drosophila. PLoS Genetics 10: e1004775.

Bergland, A. O., R. Tobler, J. González, P. Schmidt, and D. Petrov, 2016 Secondary contact and local adaptation contribute to genome-wide patterns of clinal variation in Drosophila melanogaster. Molecular Ecology 25: 1157–1174.

Bogaerts-Márquez, M., S. Guirao-Rico, M. Gautier, and J. González, 2020 Temperature, rainfall and wind variables underlie environmental adaptation in natural populations of Drosophila melanogaster. Molecular Ecology 30:938– 954.

Buels, R., E. Yao, C. M. Diesh, R. D. Hayes, M. Munoz-Torres et al., 2016 JBrowse: a dynamic web platform for genome visualization and analysis. Genome Biology 17: 66.

Burke, M. K., 2012 How does adaptation sweep through the genome? Insights from long-term selection experiments. Proceedings of the Royal Society B: Biological Sciences 279: 5029–5038.

Bushnell, B., J. Rood, and E. Singer, 2017 BBMerge – Accurate paired shotgun read merging via overlap. PLoS ONE 12: e0185056.

Campo, D., K. Lehmann, C. Fjeldsted, T. Souaiaia, J. Kao et al., 2013 Whole-genome sequencing of two North American Drosophila melanogaster populations reveals genetic differentiation and positive selection. Molecular Ecology 22: 5084–5097.

Capy, P., and P. Gibert, 2004 Drosophila melanogaster, Drosophila simulans: so similar yet so different. Genetica: 120(1-3): 5–16.

Caracristi, G., and C. Schlötterer, 2003 Genetic Differentiation Between American and European Drosophila melanogaster Populations Could Be Attributed to Admixture of African Alleles. Molecular Biology and Evolution 20: 792–799.

Celniker, S. E., and G. M. Rubin, 2003 The Drosophila melanogaster genome. Annual Review of Genomics Human Genetics 4: 89–117.

Cheng, C., B. J. White, C. Kamdem, K. Mockaitis, C. Costantini et al., 2012 Ecological Genomics of Anopheles gambiae Along a Latitudinal Cline: A Population-Resequencing Approach. Genetics 190: 1417–1432.

Cingolani, P., A. Platts, L. Wang, M. Coon, T. Nguyen et al., 2012 A program for annotating and predicting the effects of single nucleotide polymorphisms, SnpEff. Fly 6: 80–92.

Comeron, J. M., R. Ratnappan, and S. Bailin, 2012 The Many Landscapes of Recombination in Drosophila melanogaster. PLoS Genetics 8: e1002905.

Corbett-Detig, R., and R. Nielsen, 2017 A Hidden Markov Model Approach for Simultaneously Estimating Local Ancestry and Admixture Time Using Next Generation Sequence Data in Samples of Arbitrary Ploidy. PLoS Genet 13: e1006529.

Danecek, P., A. Auton, G. Abecasis, C. A. Albers, E. Banks et al., 2011 The variant call format and VCFtools. Bioinformatics 27: 2156–2158.

David, J. R., and P. Capy, 1988 Genetic variation of Drosophila melanogaster natural populations. Trends in Genetics 4: 106–111.

Deitz, K. C., G. A. Athrey, M. Jawara, H. J. Overgaard, A. Matias et al., 2016 Genome-Wide Divergence in the West-African Malaria Vector Anopheles melas. G3: Genes, Genomes, Genetics 6: 2867–2879.

Duchen, P., D. Živković, S. Hutter, W. Stephan, and S. Laurent, 2013 Demographic Inference Reveals African and European Admixture in the North American Drosophila melanogaster Population. Genetics 193: 291–301.

Duffy, J. B., 2002 GAL4 system in Drosophila: a fly geneticist’s Swiss army knife. Genesis 34: 1–15.

Durmaz, E., C. Benson, M. Kapun, P. Schmidt, and T. Flatt, 2018 An inversion supergene in Drosophila underpins latitudinal clines in survival traits. Journal of Evolutionary Biology 31: 1354–1364.

Durmaz E, Kerdaffrec E, Katsianis G, Kapun M, Flatt T. 2020. How Selection Acts on Chromosomal Inversions. In: eLS. American Cancer Society. p. 307–315.

Durmaz, E., S. Rajpurohit, N. Betancourt, D. K. Fabian, M. Kapun et al., 2019 A clinal polymorphism in the insulin signaling transcription factor foxo contributes to life-history adaptation in Drosophila. Evolution 73:1774–1792.

Erickson, P. A., C. A. Weller, D. Y. Song, A. S. Bangerter, P. Schmidt et al., 2020 Unique genetic signatures of local adaptation over space and time for diapause, an ecologically relevant complex trait, in Drosophila melanogaster. PLoS Genetics.: 16(11): e1009110.

Fabian, D. K., M. Kapun, V. Nolte, R. Kofler, P. S. Schmidt et al., 2012 Genome-wide patterns of latitudinal differentiation among populations of Drosophila melanogaster from North America. Molecular Ecology 21: 4748–4769.

Feder, A. F., D. A. Petrov, and A. O. Bergland, 2012 LDx: Estimation of Linkage Disequilibrium from High-Throughput Pooled Resequencing Data. PLoS ONE 7: e48588.

Ferretti, L., S. E. Ramos-Onsins, and M. Pérez-Enciso, 2013 Population genomics from pool sequencing. Molecular Ecology 22: 5561–5576.

Flatt, T., 2016 Genomics of clinal variation in Drosophila: disentangling the interactions of selection and demography. Molecular Ecology 25: 1023–1026.

Flatt, T., 2020 Life-History Evolution and the Genetics of Fitness Components in Drosophila melanogaster. Genetics 214: 3–48.

Gautier, M., J. Foucaud, K. Gharbi, T. Cézard, M. Galan et al., 2013 Estimation of population allele frequencies from next-generation sequencing data: pool-versus individual-based genotyping. Molecular Ecology 22: 3766–3779.

Giesen, A., W. U. Blanckenhorn, M. A. Schäfer, K. K. Shimizu, R. Shimizu-Inatsugi et al., 2020 Genomic signals of admixture and reinforcement between two closely related species of European sepsid flies. bioRxiv 2020.03.11.985903.

Glinka, S., L. Ometto, S. Mousset, W. Stephan, and D. Lorenzo, 2003 Demography and natural selection have shaped genetic variation in Drosophila melanogaster: a multi-locus approach. Genetics 165: 1269–1278.

Gould, B. A., Y. Chen, and D. B. Lowry, 2017 Pooled Ecotype Sequencing Reveals Candidate Genetic Mechanisms for Adaptive Differentiation and Reproductive Isolation. Molecular Ecology 26: 163–177.

Grenier, J. K., J. R. Arguello, M. C. Moreira, S. Gottipati, J. Mohammed et al., 2015 Global Diversity Lines–A Five-Continent Reference Panel of Sequenced Drosophila melanogaster Strains. G3: Genes, Genomes, Genetics 5: 593– 603.

Guirao-Rico, S., and J. González, 2021 Benchmarking the performance of Pool-seq SNP callers using simulated and real sequencing data. Molecular Ecology Resources 21: 1216–1229.

Guirao-Rico, S., and J. González, 2019 Evolutionary insights from large scale resequencing datasets in Drosophila melanogaster. Current Opinion in Insect Science 31: 70–76.

Hales, K. G., C. A. Korey, A. M. Larracuente, and D. M. Roberts, 2015 Genetics on the Fly: A Primer on the Drosophila Model System. Genetics 201: 815–842.

Haudry, A., S. Laurent, and M. Kapun, 2020 Population genomics on the fly: recent advances in Drosophila. In: Dutheil J.Y. (eds) Statistical Population Genomics. Methods in Molecular Biology, vol 2090. Humana, New York, NY. pp. 357–396.

Hijmans, R. J., S. E. Cameron, J. L. Parra, P. G. Jones, and A. Jarvis, 2005 Very high resolution interpolated climate surfaces for global land areas. International Journal of Climatology 25: 1965–1978.

Hivert, V., R. Leblois, E. J. Petit, M. Gautier, and R. Vitalis, 2018 Measuring Genetic Differentiation from Pool-seq Data. Genetics 210: 315–330.

Hoban, S., J. L. Kelley, K. E. Lotterhos, M. F. Antolin, G. Bradburd et al., 2016 Finding the Genomic Basis of Local Adaptation: Pitfalls, Practical Solutions, and Future Directions. American Naturalist 188: 379–397.

Hu, T. T., M. B. Eisen, K. R. Thornton, and P. Andolfatto, 2013 A second-generation assembly of the Drosophila simulans genome provides new insights into patterns of lineage-specific divergence. Genome Research 23: 89–98.

Jennings, B. H., 2011 Drosophila – a versatile model in biology & medicine. Materials Today 14: 190–195.

Jombart, T., S. Devillard, and F. Balloux, 2010 Discriminant analysis of principal components: a new method for the analysis of genetically structured populations. BMC Genetics 11: 94.

de Jong, G., and Z. Bochdanovits, 2003 Latitudinal clines in Drosophila melanogaster: Body size, allozyme frequencies, inversion frequencies, and the insulin-signalling pathway. Journal of Genetics 82: 207–223.

Kao, J. Y., A. Zubair, M. P. Salomon, S. V. Nuzhdin, and D. Campo, 2015 Population genomic analysis uncovers African and European admixture in Drosophila melanogaster populations from the south-eastern United States and Caribbean Islands. Molecular Ecology 24: 1499–1509.

Kapopoulou, A., M. Kapun, B. Pieper, P. Pavlidis, R. Wilches et al., 2020 Demographic analyses of a new sample of haploid genomes from a Swedish population of Drosophila melanogaster. Scientific Reports 10: 22415.

Kapun M, Flatt T. 2019. The adaptive significance of chromosomal inversion polymorphisms in Drosophila melanogaster. Molecular Ecology 28:1263– 1282.

Kapun, M., M. G. Barrón, F. Staubach, D. J. Obbard, R. A. W. Wiberg et al., 2020 Genomic Analysis of European Drosophila melanogaster Populations Reveals Longitudinal Structure, Continent-Wide Selection, and Previously Unknown DNA Viruses. Molecular Biology and Evolution 37: 2661–2678.

Kapun, M., D. K. Fabian, J. Goudet, and T. Flatt, 2016 Genomic Evidence for Adaptive Inversion Clines in Drosophila melanogaster. Molecular Biology and Evolution 33: 1317–1336.

Kapun M, Schalkwyk H van, McAllister B, Flatt T, Schlötterer C. 2014. Inference of chromosomal inversion dynamics from Pool-Seq data in natural and laboratory populations of Drosophila melanogaster. Molecular Ecology 23:1813–1827.

Keinan, A., and A. G. Clark, 2012 Recent Explosive Human Population Growth Has Resulted in an Excess of Rare Genetic Variants. Science 336: 740–743.

Keller, A., 2007 Drosophila melanogaster’s history as a human commensal. Current Biology 17: R77–R81.

Kent, W. J., A. S. Zweig, G. Barber, A. S. Hinrichs, and D. Karolchik, 2010 BigWig and BigBed: enabling browsing of large distributed datasets. Bioinformatics 26: 2204–2207.

Koboldt, D. C., K. Chen, T. Wylie, D. E. Larson, M. D. McLellan et al., 2009 VarScan: variant detection in massively parallel sequencing of individual and pooled samples. Bioinformatics 25: 2283–2285.

Koboldt, D. C., Q. Zhang, D. E. Larson, D. Shen, M. D. McLellan et al., 2012 VarScan 2: Somatic mutation and copy number alteration discovery in cancer by exome sequencing. Genome Research 22: 568–576.

Kofler, R., P. Orozco-terWengel, N. De Maio, R. V. Pandey, V. Nolte et al., 2011a PoPoolation: A Toolbox for Population Genetic Analysis of Next Generation Sequencing Data from Pooled Individuals. PLoS ONE 6: e15925.

Kofler, R., R. V. Pandey, and C. Schlotterer, 2011b PoPoolation2: identifying differentiation between populations using sequencing of pooled DNA samples (Pool-Seq). Bioinformatics 27: 3435–3436.

Kofler, R., and C. Schlötterer, 2014 A guide for the design of evolve and resequencing studies. Molecular Biology and Evolution 31: 474–483.

Kolaczkowski, B., A. D. Kern, A. K. Holloway, and D. J. Begun, 2011 Genomic Differentiation Between Temperate and Tropical Australian Populations of Drosophila melanogaster. Genetics 187: 245–260.

Kreitman, M., 1983 Nucleotide polymorphism at the alcohol dehydrogenase locus of Drosophila melanogaster. Nature 304: 412–417.

Kuhn, R. M., D. Haussler, and W. J. Kent, 2013 The UCSC genome browser and associated tools. Briefings in Bioinformatics 14: 144–161.

Lachaise, D., M.-L. Cariou, J. R. David, F. Lemeunier, L. Tsacas et al., 1988 Historical Biogeography of the Drosophila melanogaster Species Subgroup, pp. 159–225 in Evolutionary Biology, edited by M. K. Hecht, B. Wallace, and G. T. Prance. Springer US, Boston, MA.

Lack, J. B., C. M. Cardeno, M. W. Crepeau, W. Taylor, R. B. Corbett-Detig et al., 2015 The Drosophila Genome Nexus: A Population Genomic Resource of 623 Drosophila melanogaster Genomes, Including 197 from a Single Ancestral Range Population. Genetics 199: 1229–1241.

Lack, J. B., J. D. Lange, A. D. Tang, C.-D. B Russell, and J. E. Pool, 2016 A Thousand Fly Genomes: An Expanded Drosophila Genome Nexus. Molecular Biology and Evolution 33: 3308–3313.

Langley, C. H., K. Stevens, C. Cardeno, Y. C. G. Lee, D. R. Schrider et al., 2012 Genomic variation in natural populations of Drosophila melanogaster. Genetics 192: 533–598.

Larracuente, A. M., and D. M. Roberts, 2015 Genetics on the Fly: A Primer on the Drosophila Model System. Genetics 201: 815–842.

Lemeunier F, Aulard S, 1992. Inversion polymorphism in Drosophila melanogaster. In: Krimbas CB, Powell JR, editors. CRC Press, CRC Press. p. 576.

Li, H., 2013 Aligning sequence reads, clone sequences and assembly contigs with BWA-MEM. 1303.3997.

Li, H., 2011 Tabix: fast retrieval of sequence features from generic TAB-delimited files. Bioinformatics 27: 718–719.

Li, H., B. Handsaker, A. Wysoker, T. Fennell, J. Ruan et al., 2009 The Sequence Alignment/Map format and SAMtools. Bioinformatics 25: 2078–2079.

Li, H., and W. Stephan, 2006 Inferring the Demographic History and Rate of Adaptive Substitution in Drosophila. PLoS Genet 2: 10.

Liao, J. J. Z., and J. W. Lewis, 2000 A Note on Concordance Correlation Coefficient. FjPDA Journal of Pharmaceutical Science and Technology 54: 23–26.

Lin, L. I.-K., 1989 A Concordance Correlation Coefficient to Evaluate Reproducibility. Biometrics 45: 255–268.

Lynch, M., D. Bost, S. Wilson, T. Maruki, and S. Harrison, 2014 Population-Genetic Inference from Pooled-Sequencing Data. Genome Biology and Evolution 6: 1210–1218.

Machado, H. E., A. O. Bergland, K. R. O’Brien, E. L. Behrman, P. S. Schmidt et al., 2016 Comparative population genomics of latitudinal variation in Drosophila simulans and Drosophila melanogaster. Molecular Ecology 25: 723–740.

Machado, H. E., A. O. Bergland, R. Taylor, S. Tilk, E. Behrman et al., 2021 Broad geographic sampling reveals predictable, pervasive, and strong seasonal adaptation in Drosophila. bioRxiv 337543.

Mackay, T. F. C., S. Richards, E. A. Stone, A. Barbadilla, J. F. Ayroles et al., 2012 The Drosophila melanogaster Genetic Reference Panel. Nature 482: 173– 178.

Markow, T. A., and P. M. O’Grady, 2006 Drosophila: a guide to species identification and use. Elsevier/Academic Press Amsterdam, Boston.

Martin, M., 2011 Cutadapt removes adapter sequences from high-throughput sequencing reads. EMBnet.journal 17: 10–12.

Mateo, L., G. E. Rech, and J. González, 2018 Genome-wide patterns of local adaptation in Western European Drosophila melanogaster natural populations. Scientific Reports 8: 16143.

Mateo, L., A. Ullastres, and J. González, 2014 A transposable element insertion confers xenobiotic resistance in Drosophila. PLoS Genetics 10: e1004560.

McKenna, A., M. Hanna, E. Banks, A. Sivachenko, K. Cibulskis et al., 2010 The Genome Analysis Toolkit: A MapReduce framework for analyzing next-generation DNA sequencing data. Genome Research 20: 1297–1303.

Mölder, F., K.P. Jablonski, B. Letcher, M.B. Hall, C.H. Tomkins-Tinch, et al, 2021. Sustainable data analysis with Snakemake. F1000Res 10, 33.

Novembre, J., and M. Stephens, 2008 Interpreting principal component analyses of spatial population genetic variation. Nature Genetics 40: 646–649.

Nunez, J.C.B., Paris M, Machado, H., Bogaerts, M., Gonzalez, J., Flatt, T., Coronado, M., Kapun, M., Schmidt, P., Petrov, D., et al 2021. Note: Updating the metadata of four misidentified samples in the DrosRTEC dataset. bioRxiv:2021.01.26.428249.

Ometto, L., 2010 Inferring the Effects of Demography and Selection on Drosophila melanogaster Populations from a Chromosome-Wide Scan of DNA Variation. Molecular Biology and Evolution 22: 2119–2130.

Orozco-terWengel, P., M. Kapun, V. Nolte, R. Kofler, T. Flatt et al., 2012 Adaptation of Drosophila to a novel laboratory environment reveals temporally heterogeneous trajectories of selected alleles. Molecular Ecology 21: 4931– 4941.

Paaby, A. B., A. O. Bergland, E. L. Behrman, and P. S. Schmidt, 2014 A highly pleiotropic amino acid polymorphism in the Drosophila insulin receptor contributes to life-history adaptation. Evolution 68: 3395–3409.

Paaby, A. B., M. J. Blacket, A. A. Hoffmann, and P. S. Schmidt, 2010 Identification of a candidate adaptive polymorphism for Drosophila life history by parallel independent clines on two continents. Molecular Ecology 19: 760–774.

Pool, J. E. and Aquadro, C, 2006. History and Structure of Sub-Saharan Populations of Drosophila melanogaster. Genetics 174: 915–929.

Pool, J. E., 2015 The Mosaic Ancestry of the Drosophila Genetic Reference Panel and the D. melanogaster Reference Genome Reveals a Network of Epistatic Fitness Interactions. Molecular Biology and Evolution 32: 3236–3251.

Pool, J. E., C.-D. B Russell, R. P. Sugino, K. A. Stevens, C. M. Cardeno et al., 2012 Population Genomics of Sub-Saharan Drosophila melanogaster: African Diversity and Non-African Admixture. PLoS Genetics 8: e1003080.

Raineri, E., L. Ferretti, E.-C. Anna, B. Nevado, S. Heath et al., 2012 SNP calling by sequencing pooled samples. BMC Bioinformatics 13: 1–8.

Ramaekers, A., A. Claeys, M. Kapun, E. Mouchel-Vielh, D. Potier et al., 2019 Altering the Temporal Regulation of One Transcription Factor Drives Evolutionary Trade-Offs between Head Sensory Organs. Developmental Cell 50: 780–792.

Reinhardt, J., B. Kolaczkowski, C. Jones, D. Begun, and A. Kern, 2014 Parallel Geographic Variation in Drosophila melanogaster. Genetics 197: 361–373.

Rodríguez-Trelles, F., R. Tarrío, and M. Santos, 2013 Genome-wide evolutionary response to a heat wave in Drosophila. Biology Letters 9: 20130228.

Rudman, S. M., S. Greenblum, R. C. Hughes, S. Rajpurohit, O. Kiratli et al., 2019 Microbiome composition shapes rapid genomic adaptation of Drosophila melanogaster. FProceedings of the National Academy of Sciences USA 116: 20025–20032.

dos Santos, G., A. J. Schroeder, J. L. Goodman, V. B. Strelets, M. A. Crosby et al., 2015 FlyBase: introduction of the Drosophila melanogaster Release 6 reference genome assembly and large-scale migration of genome annotations. Nucleic Acids Research 43: D690–D697.

Schlötterer, C., R. Tobler, R. Kofler, and V. Nolte, 2014 Sequencing pools of individuals — mining genome-wide polymorphism data without big funding. Nature Reviews Genetics 15: 749–763.

Schneider, D., 2000 Using Drosophila as a model insect. Nature Reviews Genetics 1: 218–226.

Sprengelmeyer, Q. D., S. Mansourian, J. D. Lange, D. R. Matute, B. S. Cooper et al., 2020 Recurrent Collection of Drosophila melanogaster from Wild African Environments and Genomic Insights into Species History. Molecular Biology and Evolution 37: 627–638.

Stajich, J. E., and M. W. Hahn, 2005 Disentangling the Effects of Demography and Selection in Human History. Molecular Biology and Evolution 22: 63–73.

Stephan, W., 2016 Signatures of positive selection: from selective sweeps at individual loci to subtle allele frequency changes in polygenic adaptation. Molecular Ecology 25: 79–88.

Turner, T. L., M. T. Levine, M. L. Eckert, and D. J. Begun, 2008 Genomic Analysis of Adaptive Differentiation in Drosophila melanogaster. Genetics 179: 455–473.

Turner, T. L., A. D. Stewart, A. T. Fields, W. R. Rice, and A. M. Tarone, 2011 Population-Based Resequencing of Experimentally Evolved Populations Reveals the Genetic Basis of Body Size Variation in Drosophila melanogaster. PLoS Genetics 7: e1001336.

Umina, P. A., 2005 A Rapid Shift in a Classic Clinal Pattern in Drosophila Reflecting Climate Change. Science 308: 691–693.

Waldvogel, A.-M., B. Feldmeyer, G. Rolshausen, M. Exposito-Alonso, C. Rellstab et al., 2020 Evolutionary genomics can improve prediction of species’ responses to climate change. Evolution Letters 4: 4–18.

Wallace, M.A., Coffman, K.A., Gilbert, C., Ravindran, S., Albery, G.F., Abbott, J., Argyridou, E., Bellosta, P., Betancourt, A.J., Colinet, H., et al., 2021 The discovery, distribution and diversity of DNA viruses associated with Drosophila melanogaster in Europe. Virus Evolution. veab031.

Wittmann, M. J., A. O. Bergland, M. W. Feldman, P. S. Schmidt, and D. A. Petrov Seasonally fluctuating selection can maintain polymorphism at many loci via segregation lift. Proceedings of the National Academy of Sciences USA 114: E9932–E9941.

Wright, S., 1943 Isolation by distance. Genetics 28: 114.

Zheng, X., S. M. Gogarten, M. Lawrence, A. Stilp, M. P. Conomos et al., 2017 SeqArray—a storage-efficient high-performance data format for WGS variant calls. Bioinformatics 33: 2251–2257.

Zhu, Y., A. O. Bergland, J. González, and D. A. Petrov, 2012 Empirical Validation of Pooled Whole Genome Population Re-Sequencing in Drosophila melanogaster. PLoS ONE 7: e41901.

