## Supplemental Material for "*Drosophila* Evolution over Space and Time (DEST) - A New Population Genomics Resource"

### Supplemental Material, Tables

**Supplemental Material, Table S1.** Population collection information and basic meta-data.

**Supplemental Material, Table S2.** Basic library statistics for each population.

**Supplemental Material, Table S3.** $p_N/p_S$ estimates for each population based on PoolSNP and SNAPE-pooled. In addition, number of private SNPs for the SNAPE-pooled dataset.

**Supplemental Material, Table S4.** Allele frequency and SNP calling comparisons between callers.

**Supplemental Material, Table S5**. Population genetic estimates for the 271 populations used in this study. Estimates of nucleotide diversity ($\pi$), Watterson's $\theta$ and Tajima's $D$ using both the PoolSNP and SNAPE-pooled (SNAPE) datasets for each major chromosome arm (excluding the 4$^{th}$ dot chromosome).

**Supplemental Material, Figure S1:** Flowchart of mapping/analysis pipeline.

**Supplemental Material, Figure S2:** DEST browser, general description.

**Supplemental Material, Figure S3:** DEST browser, $F_{ST}$ example.

**Supplemental Material, Figure S4:** Allele frequency correlations between DEST and Kapun *et al.* (2020).

**Supplemental Material, Figure S5:** Allele frequency correlations between DEST and Machado *et al.* (2021).

**Supplemental Material, Figure S6:** Allele frequency correlations between DEST and Bergland *et al.* (2016).

**Supplemental Material, Figure S7:** Coverage correlations between DEST and Kapun *et al.* (2020).

**Supplemental Material, Figure S8:** Coverage correlations between DEST and Machado *et al.* (2021).

**Supplemental Material, Figure S9:** Coverage correlations between DEST and Bergland *et al.* (2016).

**Supplemental Material, Figure S10:** Effective coverage correlations between DEST and Kapun *et al.* (2020).

**Supplemental Material, Figure S11:** Effective coverage correlations between DEST and Machado *et al.* (2021).

**Supplemental Material, Figure S12:** Effective coverage correlations between DEST and Bergland *et al.* (2016).

**Supplemental Material, Figure S13:** Population genetic estimates for African, European and North American populations by chromosome arm. Shown are estimates of nucleotide diversity ($\pi$), Watterson's $\theta$, Tajima's $D$ and $p_N/p_S$ for African, European and North American populations using both the PoolSNP and SNAPE-pooled (SNAPE) datasets for each major chromosome arm (excluding the 4$^{th}$ dot chromosome).

**Supplemental Material, Figure S14.** Biplots between PCA and latitude and longitude.

**Supplemental Material, Figure S15.** PCA of pooled vs inbreed samples. The inbred samples are as follows: Winters (CA), Ithaca (NY), and Raleigh (NC).

**Supplemental Material, Figure S16.** PCA of DEST dataset colored uniquely by continental cluster.

**Supplemental Methods.** Tutorial on how to apply DIM analysis.

**Supplemental Material, Figures**

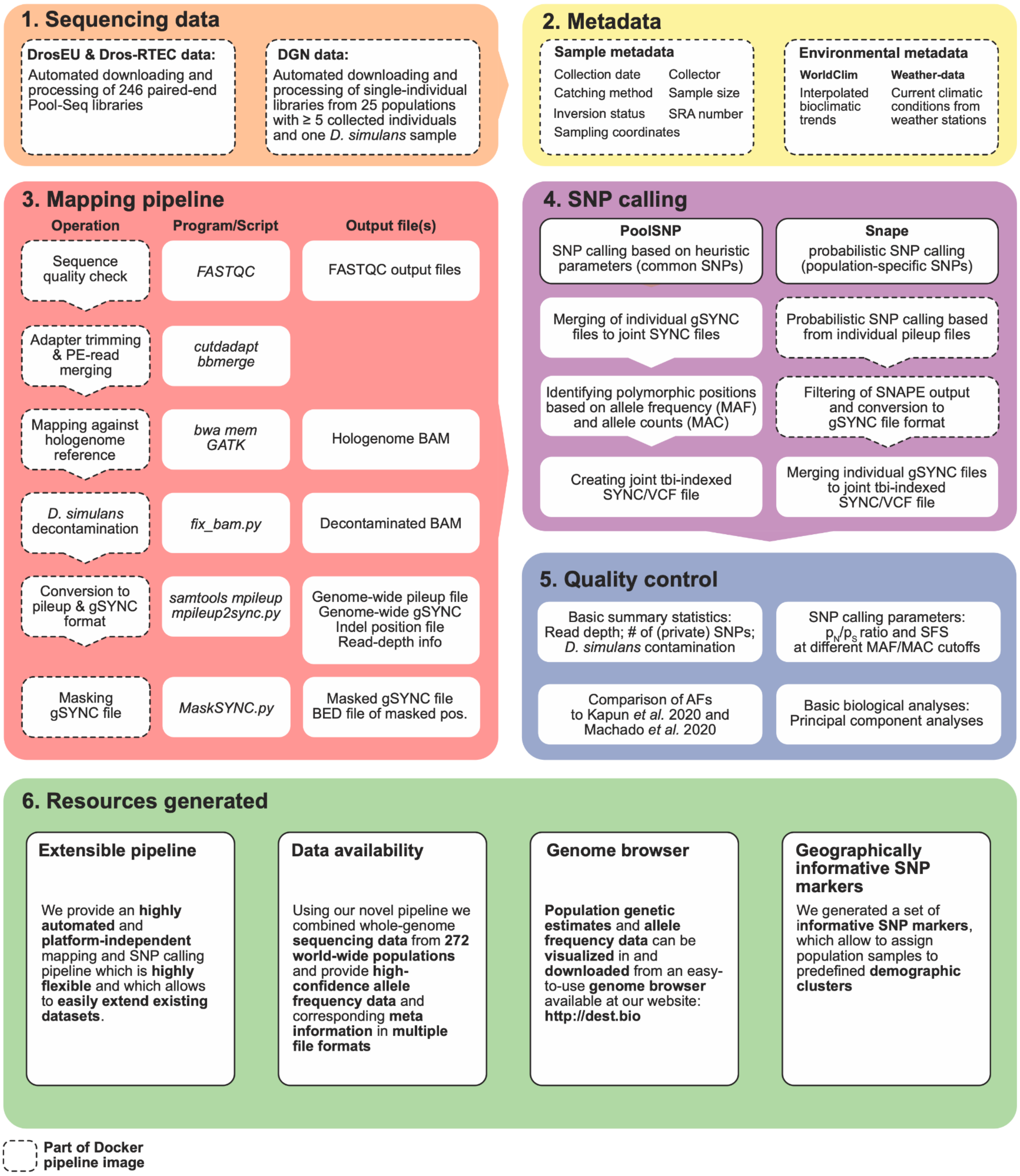


**Figure S1**. Flowchart of the bioinformatic workflow to combine and jointly analyze whole-genome sequencing data of 271 worldwide *D. melanogaster* population samples and one *D. simulans* genome, ranging from sample information (1) to resources generated in this study (6).

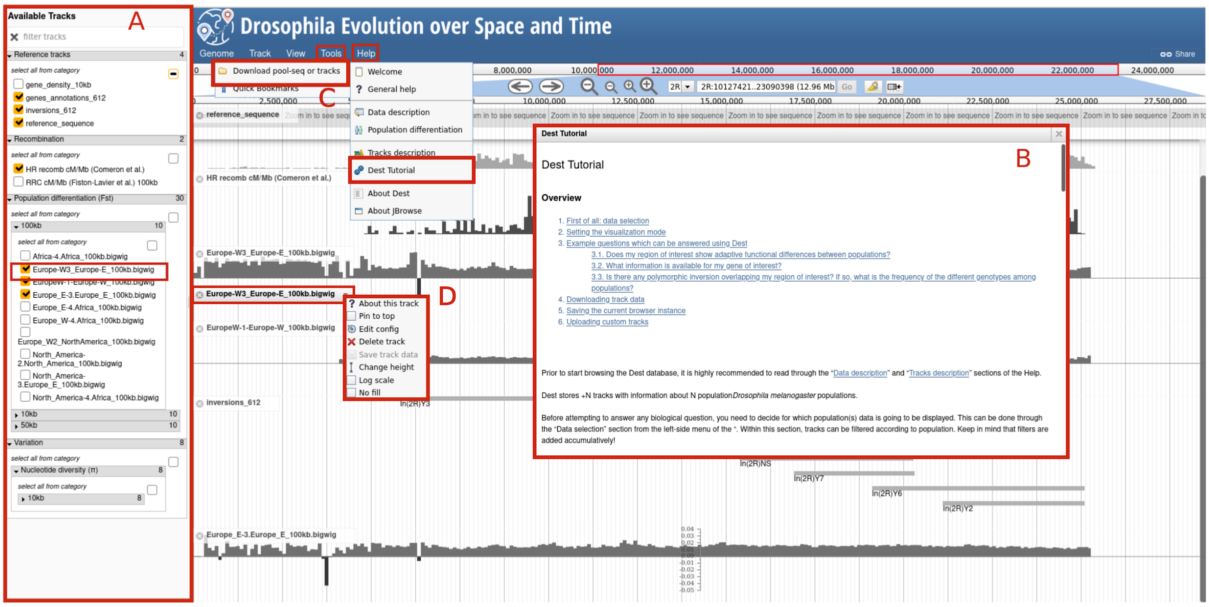


**Figure S2**. Snapshot of the Dest.bio genome browser page showing the main functionalities of the genome browser. (A) Tracks selector. (B) Tutorial. This page is accessible through the Help menu. It guides users through the processes of selecting, visualizing and downloading track data. (C) Download options. It allows downloading full track data or region data for a given set of coordinates, for one or all populations, in VCF format. (D) Track information. It shows basic information about the track and multiple options to change the default visualization.

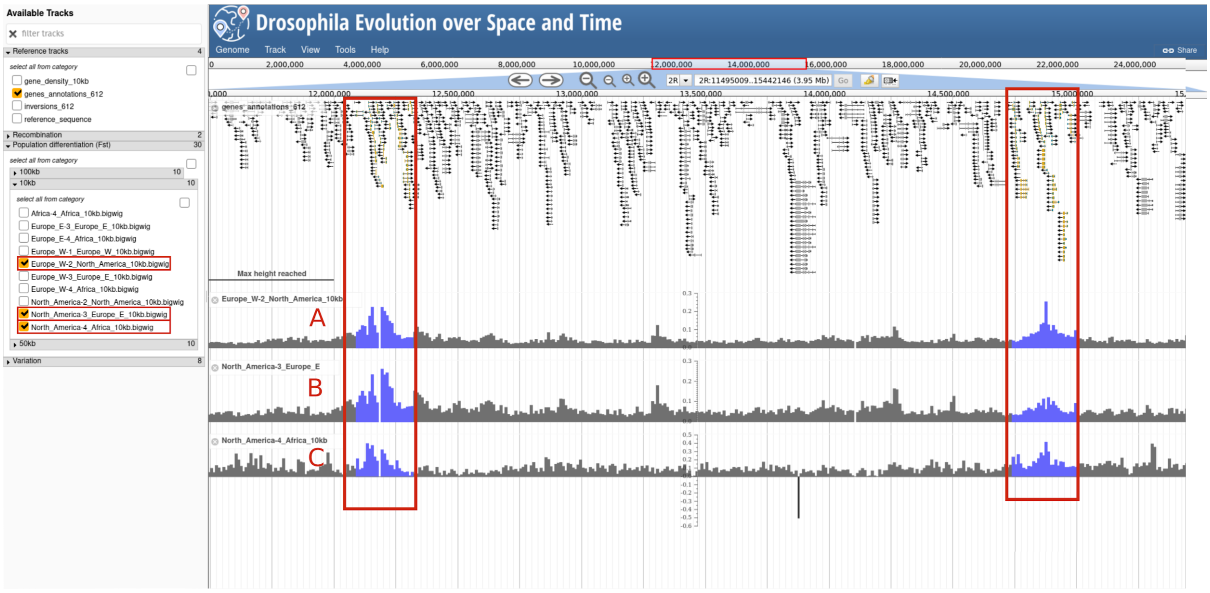


**Figure S3.** Snapshot of the dest.bio browser displaying two regions (in blue) of high population differentiation ($F_{ST}$) on chromosome arm *2R*. Both regions contain genes of the cytochrome P450 family, such as *Cyp6g1* and *Cyp6a23*, which are genes that are well-known to underlie adaptive resistance to insecticides (see Liu *et al.* 2015 for a review). A, B and C correspond to Western Europe vs. North America, Eastern Europe vs. North America and Africa vs. North America pairwise population $F_{ST}$ analyses, respectively.

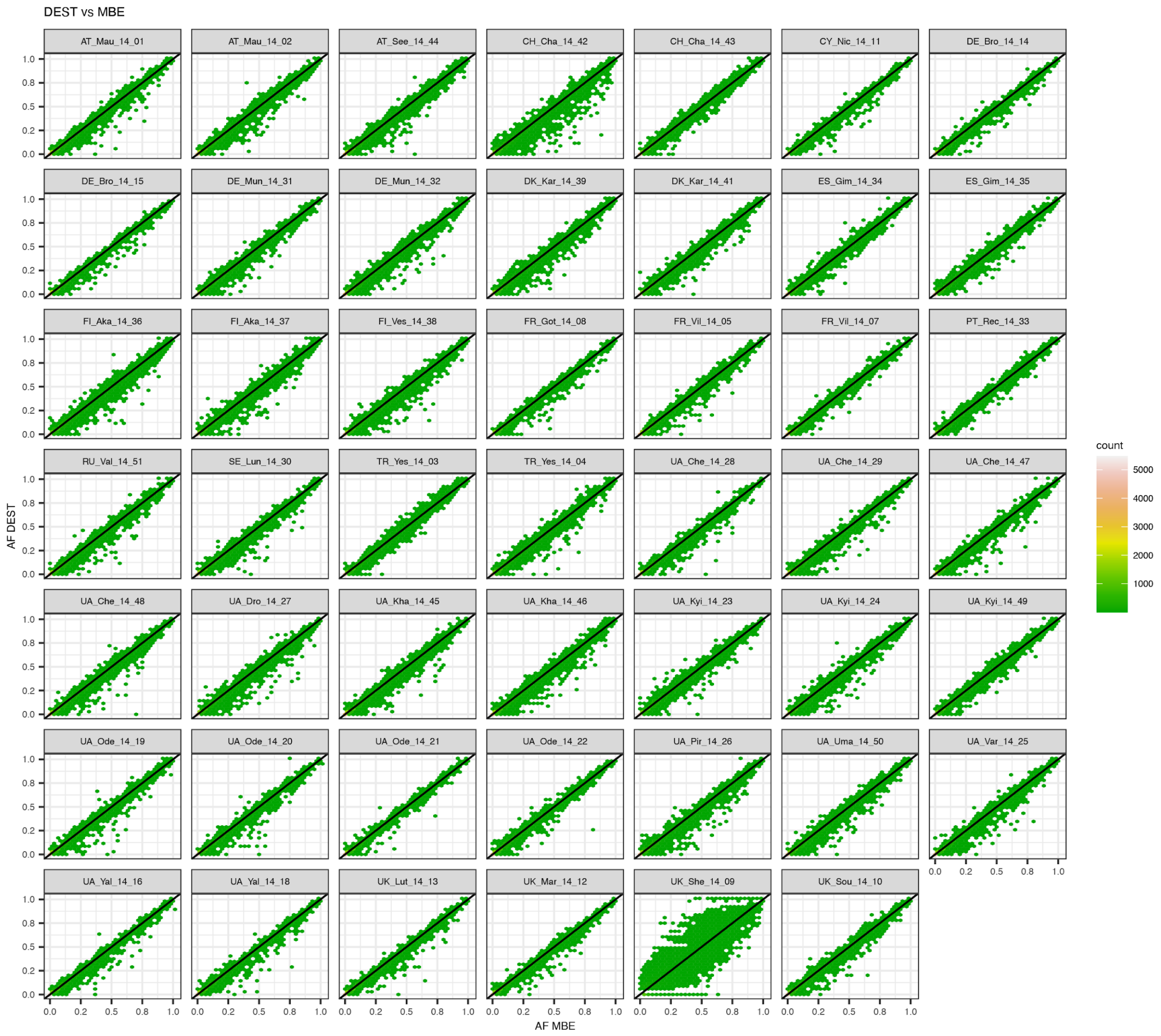


**Supplemental Material, Figure S4:** Allele frequency (AF) correlations between DEST and Kapun *et al*. (2020).

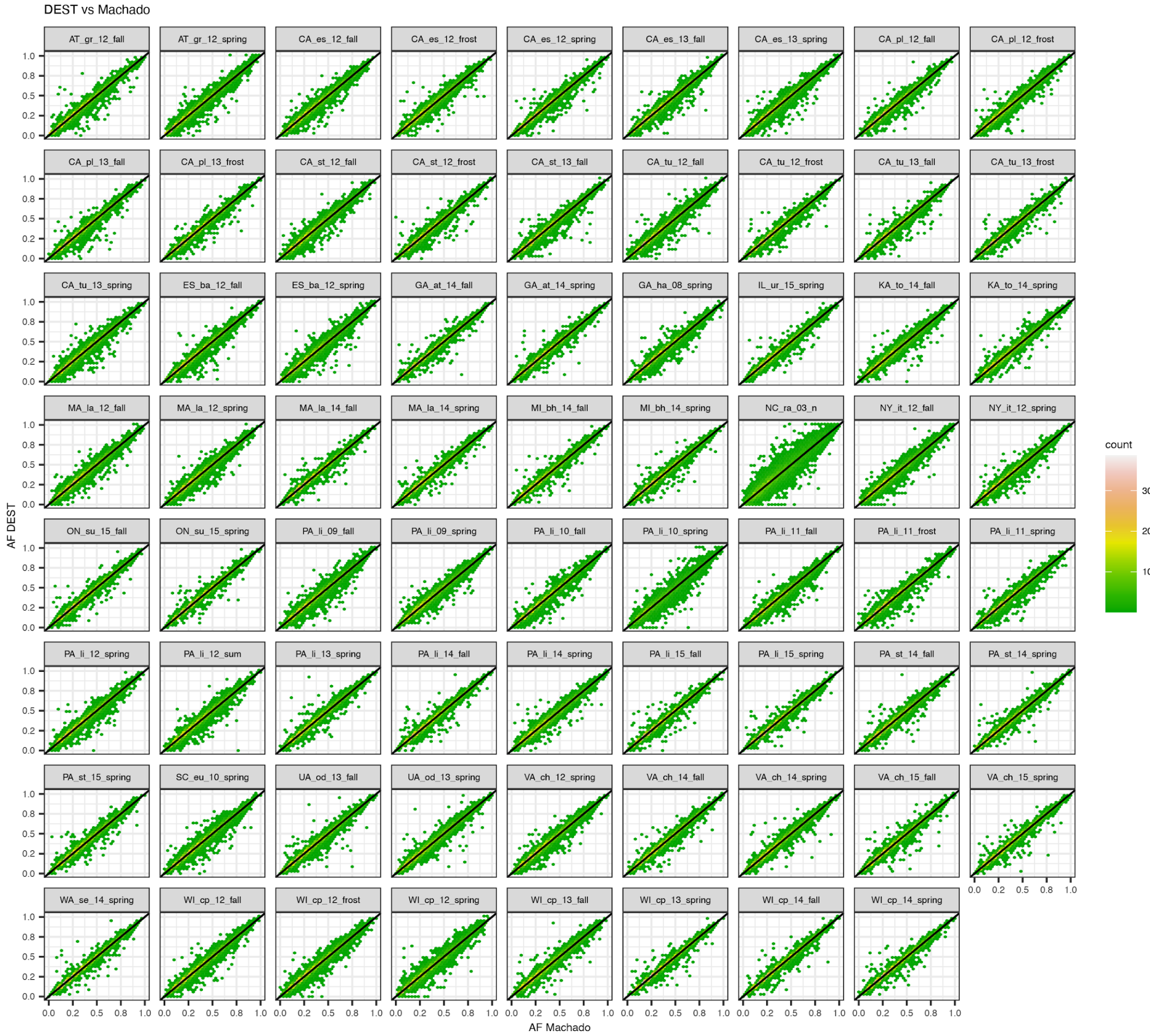


**Supplemental Material, Figure S5:** Allele frequency (AF) correlations between DEST and Machado *et al*. 2021.

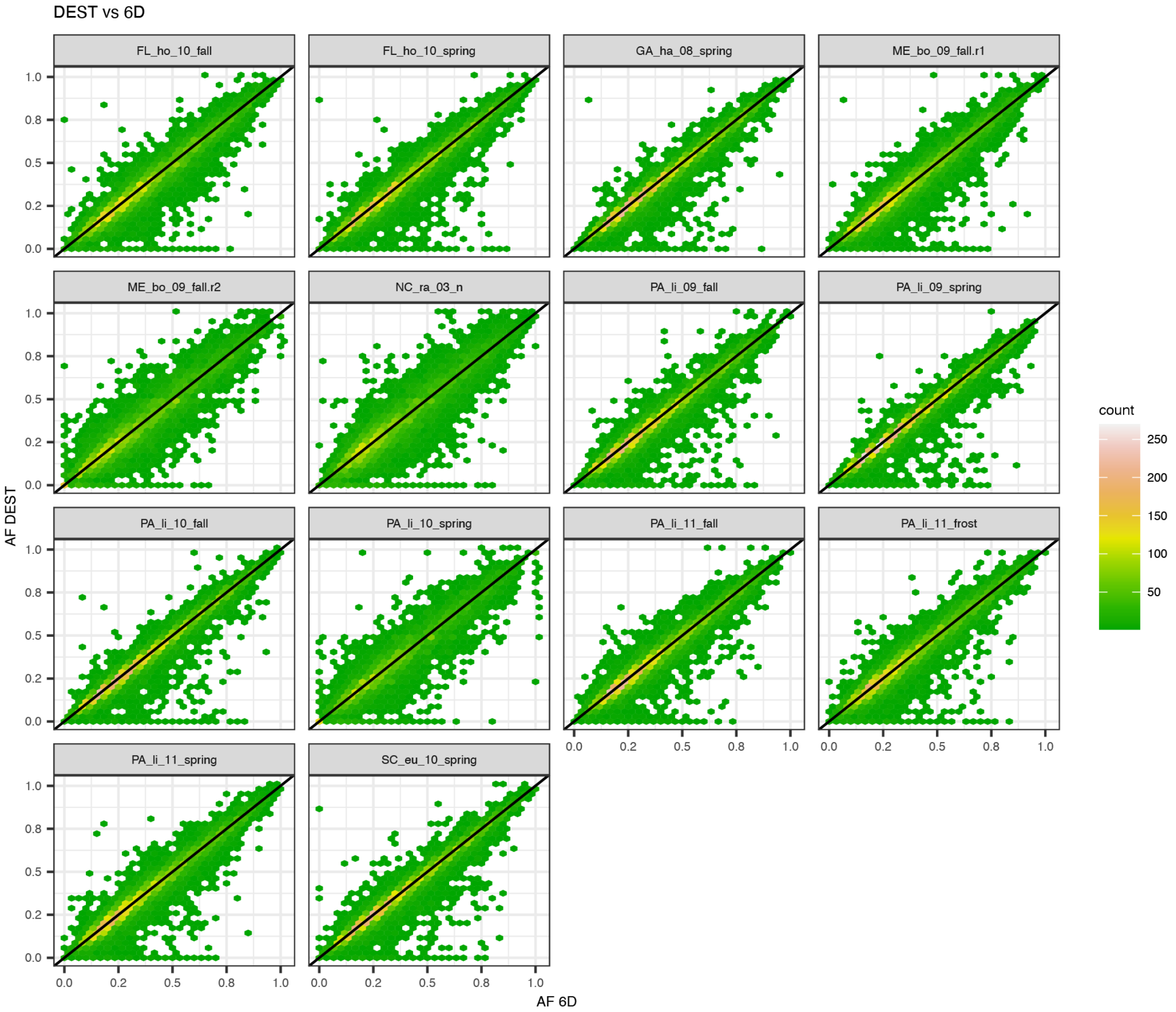


**Supplemental Material, Figure S6:** Allele frequency (AF) correlations between DEST and Bergland *et al*. (2016).

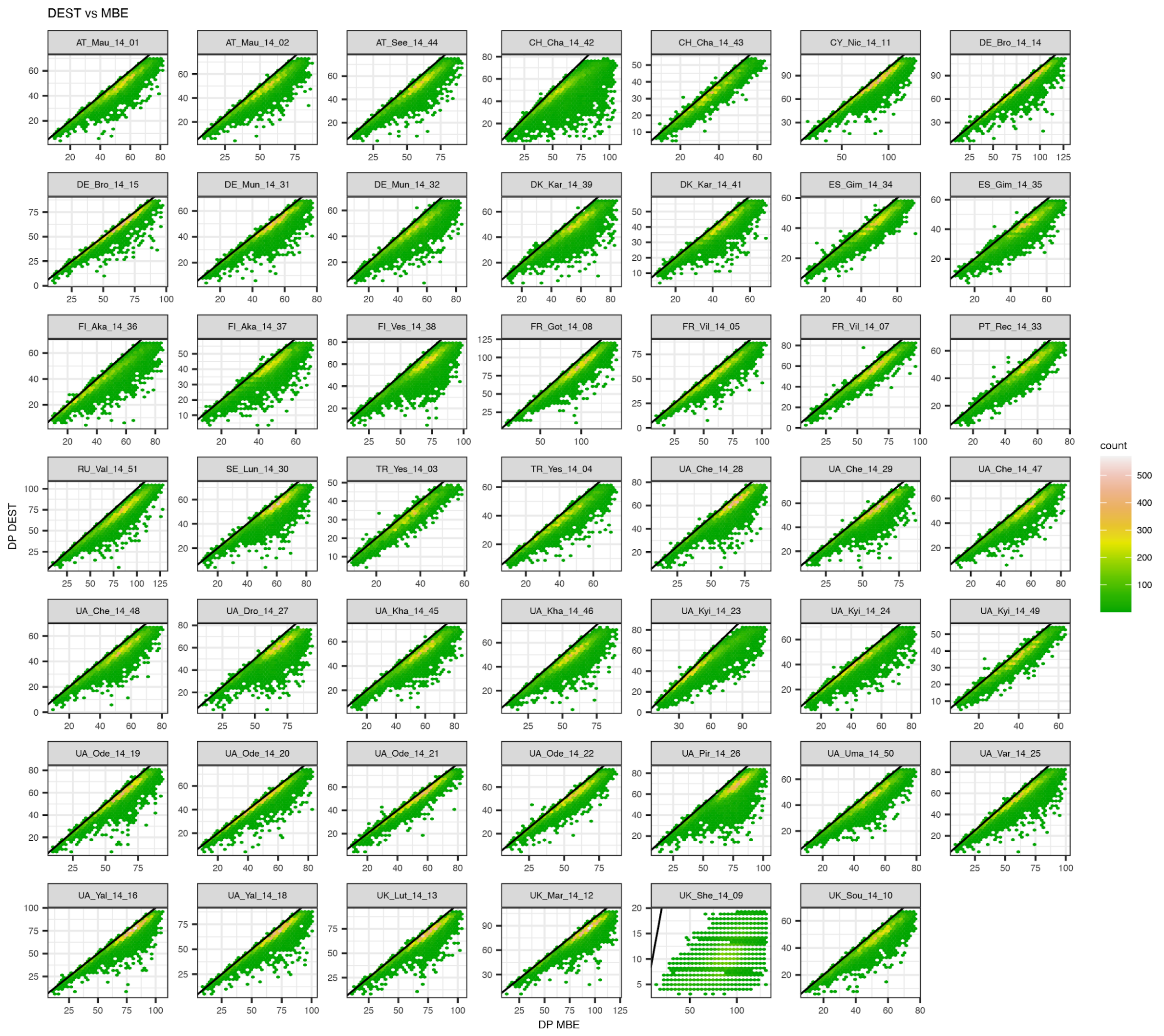


**Supplemental Material, Figure S7:** Coverage correlations between DEST and Kapun *et al*. (2020).

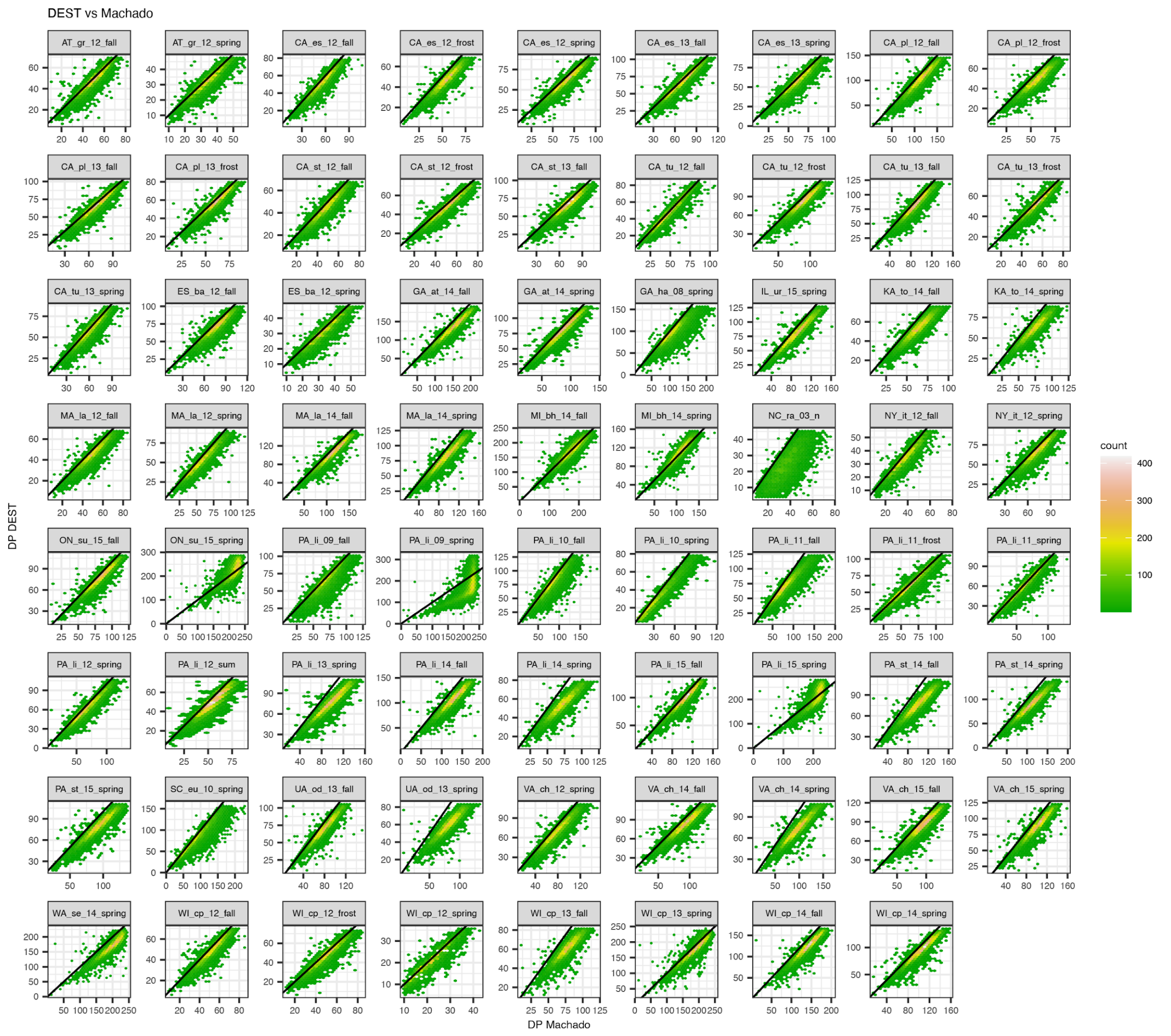


**Supplemental Material, Figure S8:** Coverage correlations between DEST and Machado *et al*. (2021).

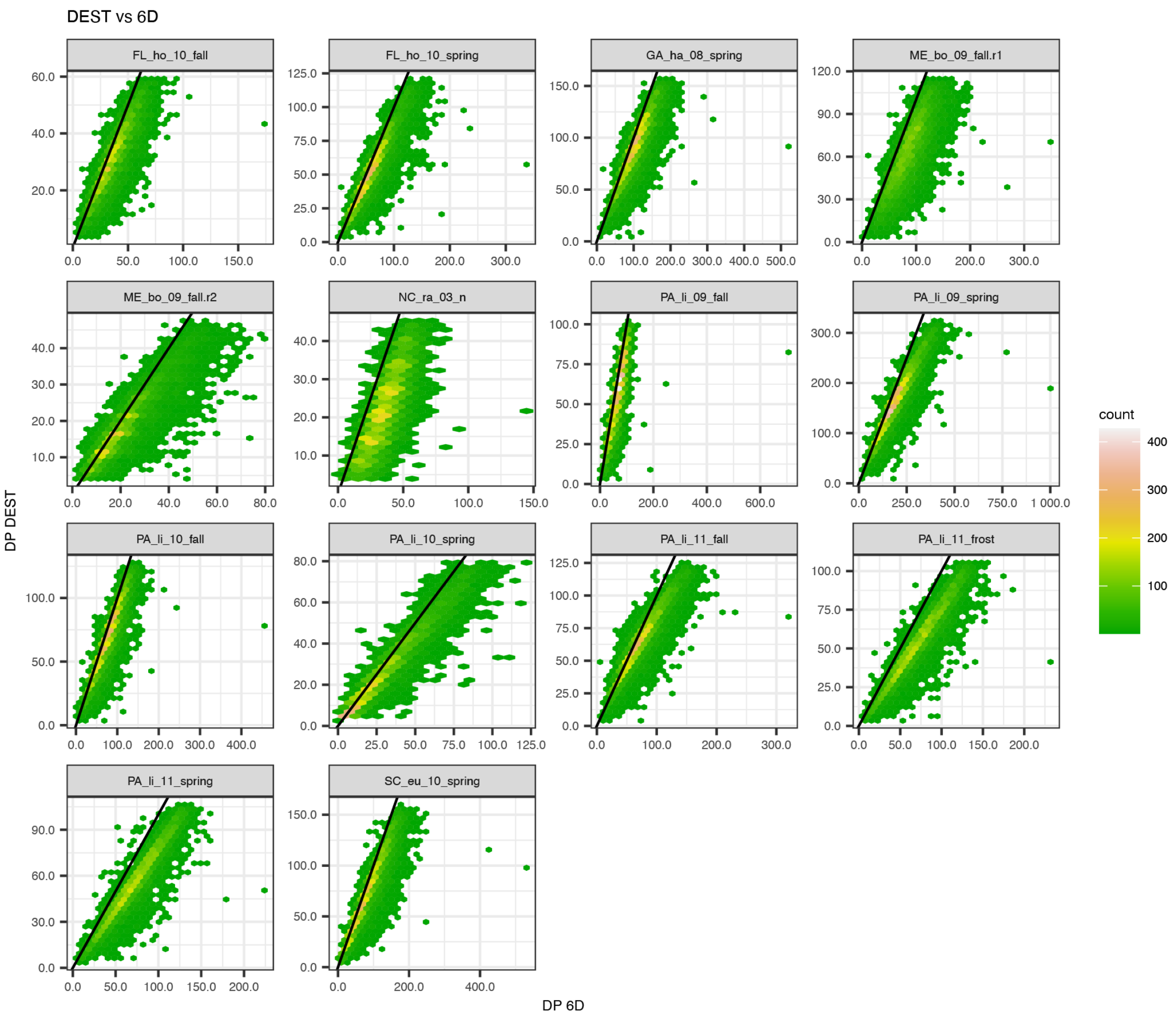


**Supplemental Material, Figure S9:** Coverage correlations between DEST and Bergland *et al*. (2016).

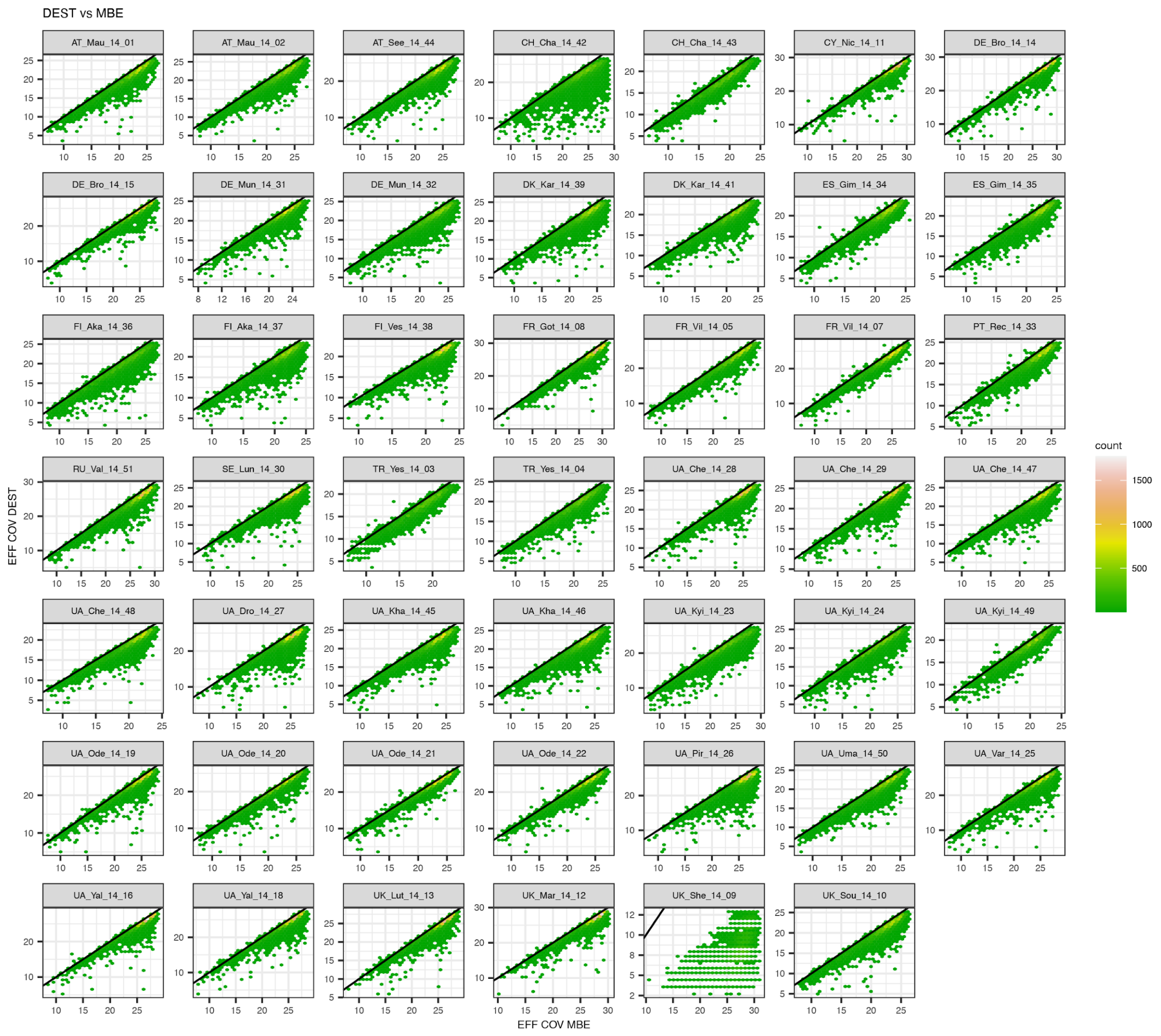


**Supplemental Material, Figure S10:** Effective coverage correlations between DEST and Kapun *et al*. (2020).

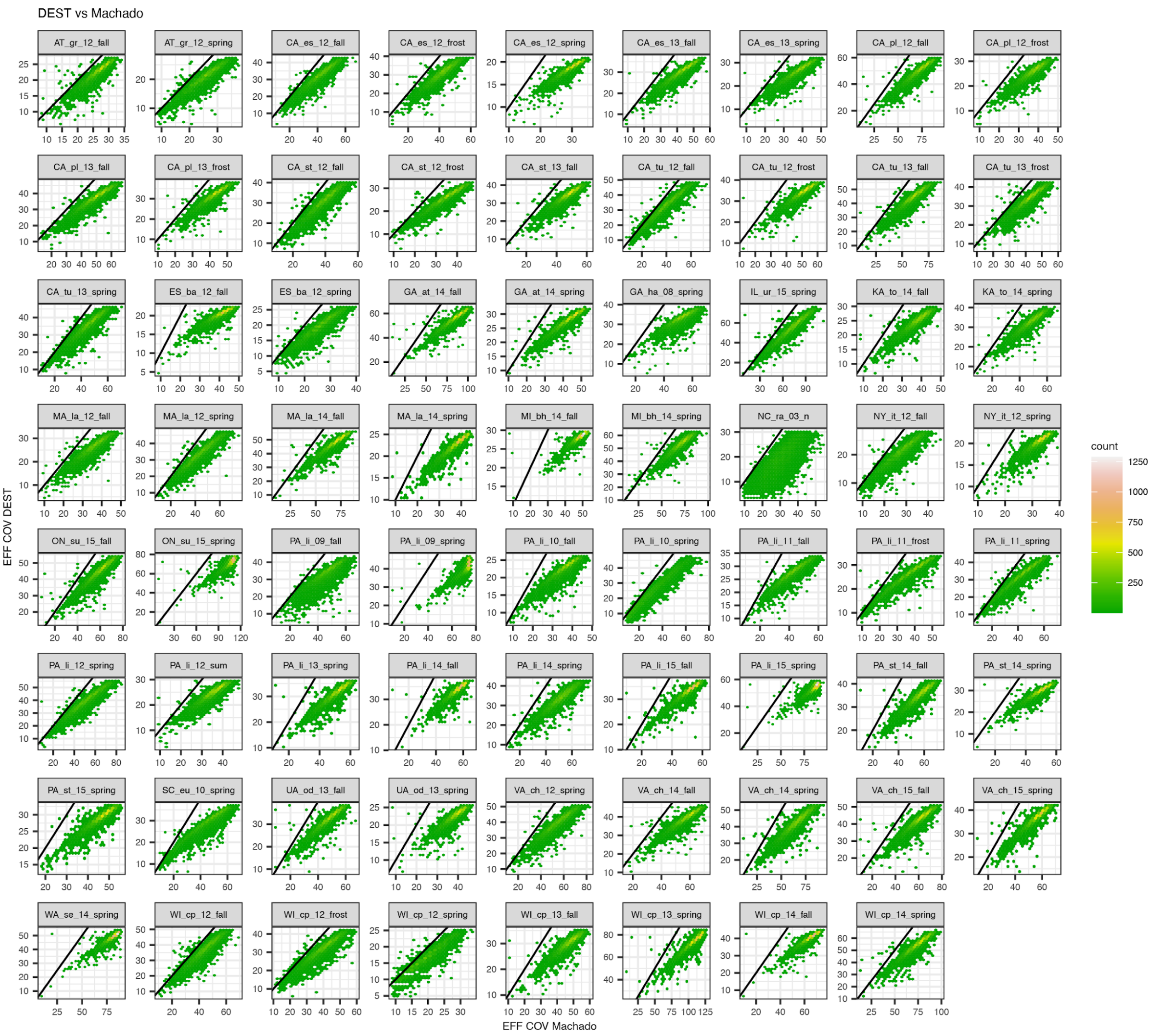


**Supplemental Material, Figure S11:** Effective coverage correlations between DEST and Machado *et al*. (2021).

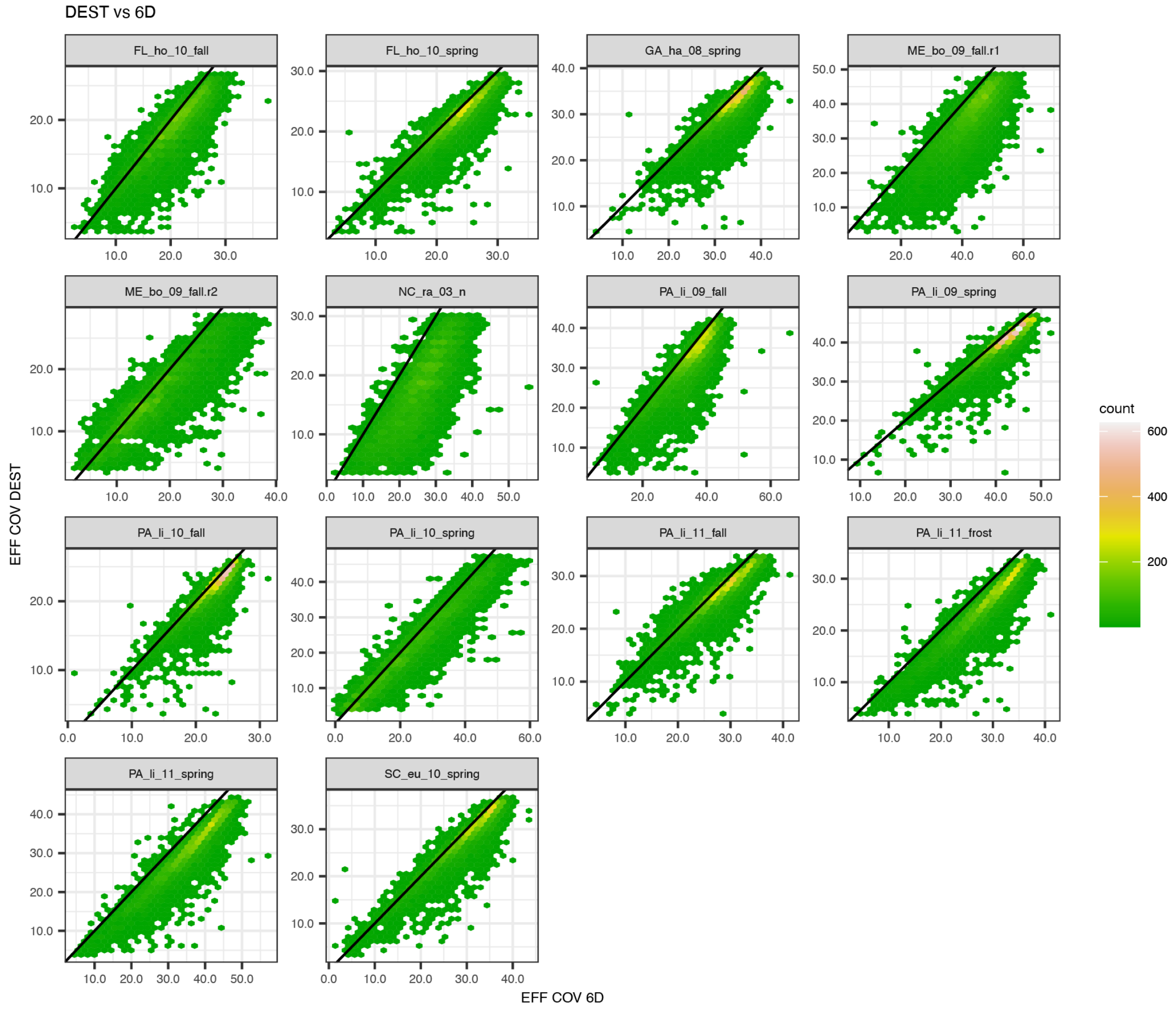


**Supplemental Material, Figure S12:** Effective coverage correlations between DEST and Bergland *et al* (2016).

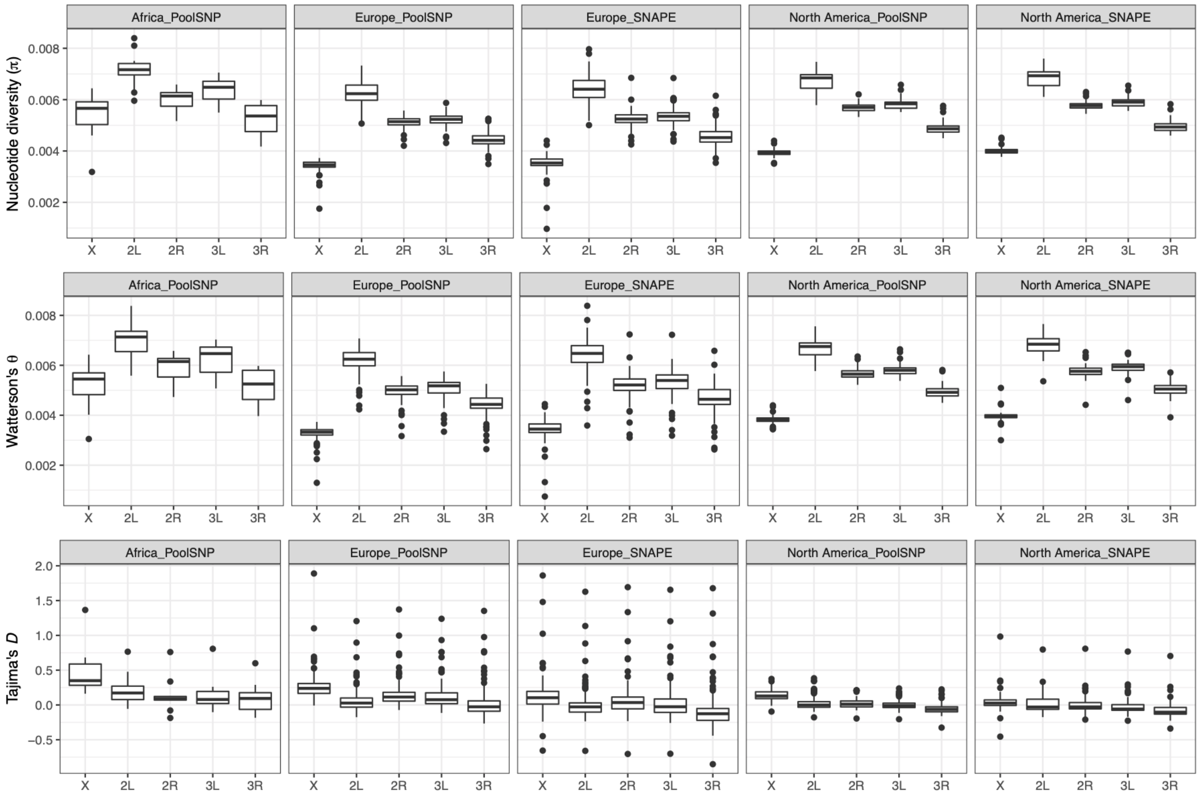


**Supplemental Material, Figure S13:** Population genetic estimates for African, European and North American populations by chromosome arm. Shown are estimates of nucleotide diversity (*π*), Watterson's *θ* and Tajima's *D* for African, European and North American populations using both the PoolSNP and SNAPE-pooled (SNAPE) datasets for each major chromosome arm (excluding the 4$^{th}$ dot chromosome).

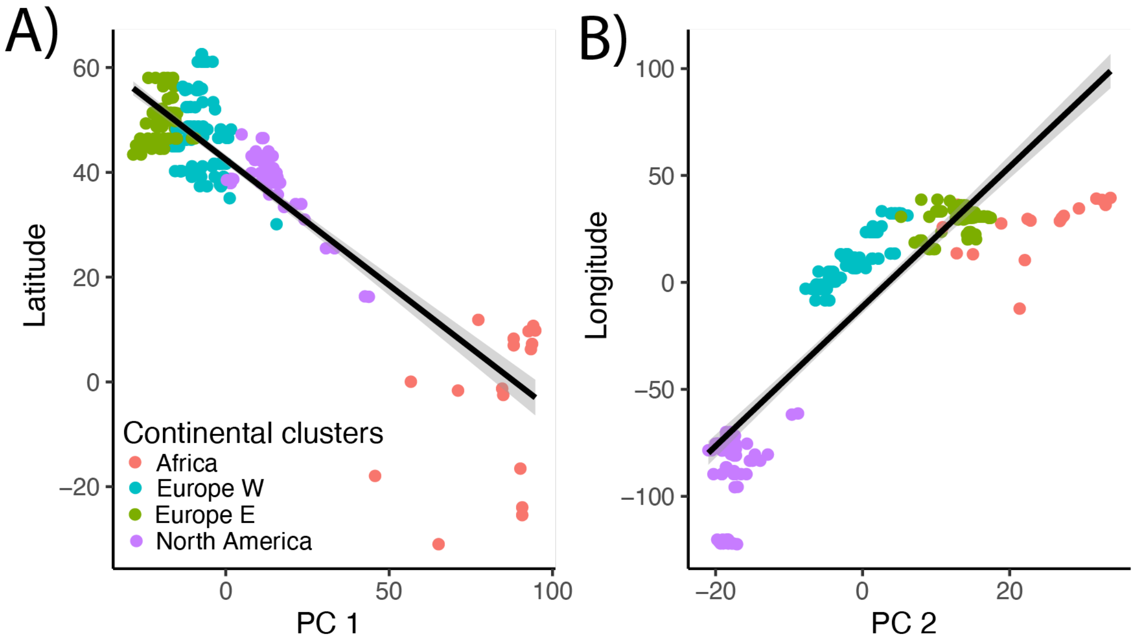


**Supplemental Material, Figure S14.** Biplots between PCA and latitude and longitude. (A) PC1, which separates African samples from all other samples, is correlated with latitude ($\rho$ = -0.86; CI = -0.88, -0.82; $p < 2.2\text{x}10^{-16}$). (B) PC2, on the other hand, divides populations and is correlated with longitude ($\rho$ = 0.87; CI = 0.84, 0.90; $p < 2.2\text{x}10^{-16}$).

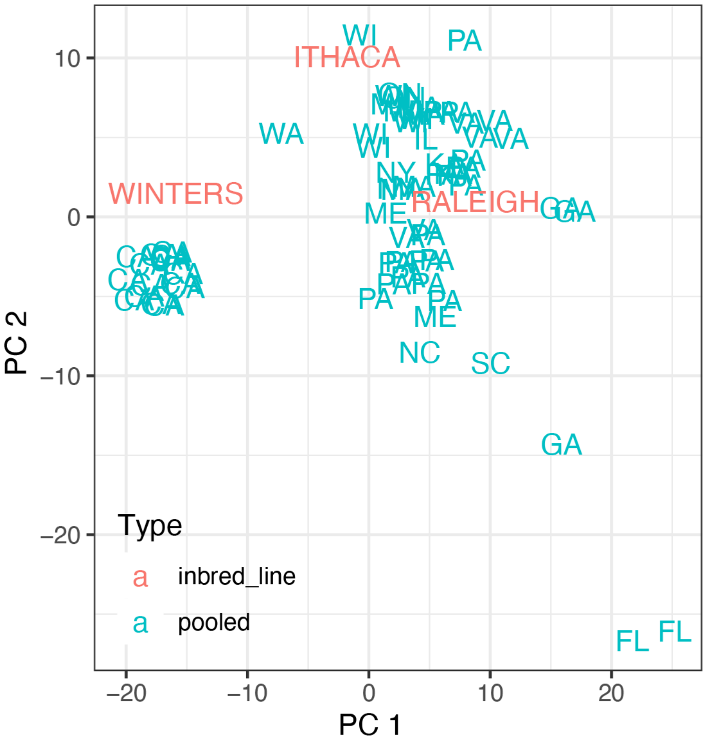


**Supplemental Material, Figure S15.** PCA of pooled vs inbred samples. The inbred samples are as follows: Winters (CA), Ithaca (NY), and Raleigh (NC).

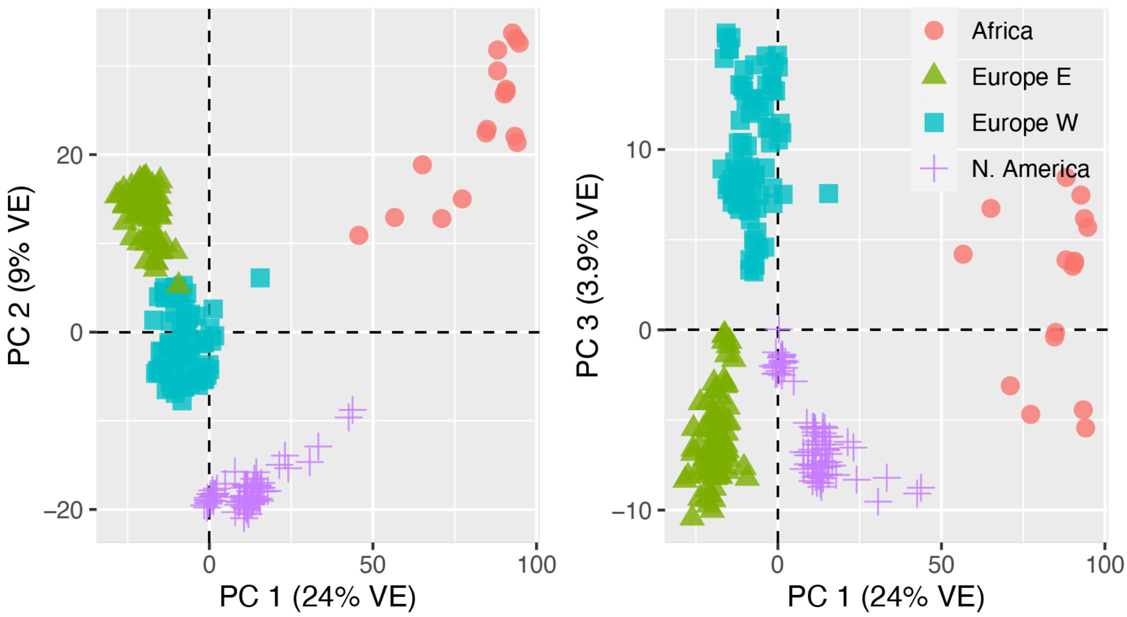


**Supplemental Material, Figure S16.** PCA of DEST dataset colored uniquely by continental cluster.
